# Sub-10 nm fluorescence imaging

**DOI:** 10.1101/2022.02.08.479592

**Authors:** Dominic A. Helmerich, Gerti Beliu, Danush Taban, Mara Meub, Marcel Streit, Alexander Kuhlemann, Sören Doose, Markus Sauer

## Abstract

Advances in superresolution microscopy demonstrated single-molecule localization precisions of a few nanometers. However, translation of such high localization precisions into sub-10 nm spatial resolution in biological samples remains challenging. Here, we show that resonance energy transfer between fluorophores separated by less than 10 nm results in accelerated fluorescence blinking and consequently lower localization probabilities impeding sub-10 nm fluorescence imaging. We demonstrate that time-resolved fluorescence detection in combination with photoswitching fingerprint analysis can be used advantageously to determine the number and distance even of spatially unresolvable fluorophores in the sub-10 nm range. In combination with genetic code expansion (GCE) with unnatural amino acids and bioorthogonal click-labeling with small fluorophores photoswitching fingerprint analysis enables sub-10 nm resolution fluorescence imaging in cells.

## Main Text

Over the past decade, superresolution fluorescence imaging by single-molecule localization has evolved as a very powerful method for subdiffraction-resolution fluorescence imaging of cells and structural investigations of cellular organelles (*1,2*). However, although single-molecule localization microscopy (SMLM) methods can now provide a spatial resolution of ~20 nm, i.e. well below the diffraction limit of light microscopy, they do not provide true molecular resolution of a few nanometers which is required to comprehensively understand the composition and 3D organization of organelles, multiprotein complexes or protein-dense networks in real biological samples such as cells or tissues.

Three central parameters that determine image resolution in SMLM experiments are the localization precision (the spread of the measured position coordinates around their mean values), the localization accuracy (the deviation of the mean of the measured coordinates from the true position), and the labeling density. Much attention focused mainly on improving the localization precision as one of the two key determinants of image resolution (*3–5*). For instance, the use of sequential structured illumination in combination with single-molecule detection as used in MINFLUX, SIMPLE and SIMFLUX allowed to improve the localization precision of *direct* stochastic optical reconstruction microscopy (*d*STORM)(*6,7*) using the red-absorbing cyanine dyes Alexa Fluor 647 (AF647) and Cy5 in photoswitching buffer to the 1-5 nm range(*8–12*). Such high localization precisions permitted to resolve some fluorophores separated by only 6 nm on DNA origami and ~10 nm in nuclear pore complexes (NPCs), respectively (*8,12*).However, the results also sparked a debate about the spatial resolution claimed and the reliability of the method (*13*). In particular, the images revealed a low detection probability of fluorophores when separated by only a few nanometers evidenced by a high number of incomplete DNA origami and missing protein localizations in the biological samples. On the other hand, these reports demonstrated that anisotropic photon emission of fluorophores due to limited rotational mobility, which has been assumed to cause substantial localization bias (*14,15*), can be neglected for highly water soluble cyanine dyes such as AF647 and Cy5. Hence, the observed low localization probability of fluorophores separated by <10 nm represents a conundrum. Since site-specific and quantitative labeling of DNA origami with fluorophores is feasible even for sub-10 nm interfluorophore distances, it remains obscured why nanometer localization precisions cannot be translated into molecular resolution with higher reliability. Or, in other words, why does the localization probability severely decrease for interfluorophore distances of <10 nm and how can we bypass this hitherto non-perceived limit? So far, a model that convincingly explains the observed behavior does not exist. To decipher the limits that SMLM methods such as *d*STORM are facing in the sub-10 nm regime and that cause the observed significant deterioration in localization probability, we studied DNA origami with different interfluorophore distances. Our data demonstrate that the on/off photoswitching kinetics strongly depend on interfluorophore distance in the sub-10 nm range. We show how photoswitching fingerprint analysis in combination with time-resolved fluorescence detection can overcome these limitations. We demonstrate the concept on DNA origami carrying four fluorophores at distances of 18, 9, 6, and 3 nm and exemplify its translation to biological systems by investigating the stoichiometry and interfluorophore distance of subunits of oligomeric receptors in cells labeled by genetic code expansion (GCE) with unnatural amino acids and click labeling using tetrazine-dyes.

### The 10 nm resolution barrier

To investigate the problems associated with sub-10 nm fluorescence imaging in more detail, we designed DNA origami (*16,17*) carrying four Cy5 dyes separated by 18, 9, 6, and 3 nm and immobilized them on coverslips via biotin-streptavidin binding (Fig. 1A and figs. S1 and S2). As reference we used the same DNA origami labeled with only a single Cy5. *d*STORM imaging was performed in standard photoswitching buffer using exclusively 640 nm irradiation. While *d*STORM can resolve the four fluorophores at 18 nm distance in some cases it cannot resolve the fluorophores separated by 9, 6, and 3 nm (Fig. 1B and fig. S3). For direct comparison we performed DNA-PAINT using Cy3B-labeled imager strands (Fig. 1C and fig. S4) (*18*). DNA-PAINT clearly achieves a higher spatial resolution but also fails to resolve the fluorophores for shorter distances of 6 and 3 nm. Even though we detected intact DNA origami carrying four fluorophores (Figs. 1B and 1C) we would like to point out that for most DNA origami investigated we could not detect four fluorophores (figs. S3 and S4). What struck us more, however, is the peculiar difference in photoswitching kinetics, i.e. blinking noticeable in the *d*STORM movies recorded from the 6 nm and 3 nm DNA origami. While the DNA-PAINT movies recorded for the different origami do not show any difference in blinking behavior throughout the entire recording time (movies S1-S5), the *d*STORM movies of the 6 nm (movie S9) and 3 nm origami (movies S10 and S11) show often a “flickering” fluorescence intensity during the first seconds, i.e. very fast blinking compared to the expected well-defined blinking of Cy5 dyes as observed for the reference (movie S6) and the 18 nm origami (movie S7). Even for the 9 nm DNA origami some fluorescent spots appear to show faster blinking (movie S8). Comparison of the fluorescence signal densities during the first seconds and after a few minutes clearly proves that faster blinking is accompanied with faster photobleaching. These observations point out that the on/off photoswitching kinetics of Cy5 dyes is substantially accelerated at shorter interfluorophore distances.

**Fig 1.**
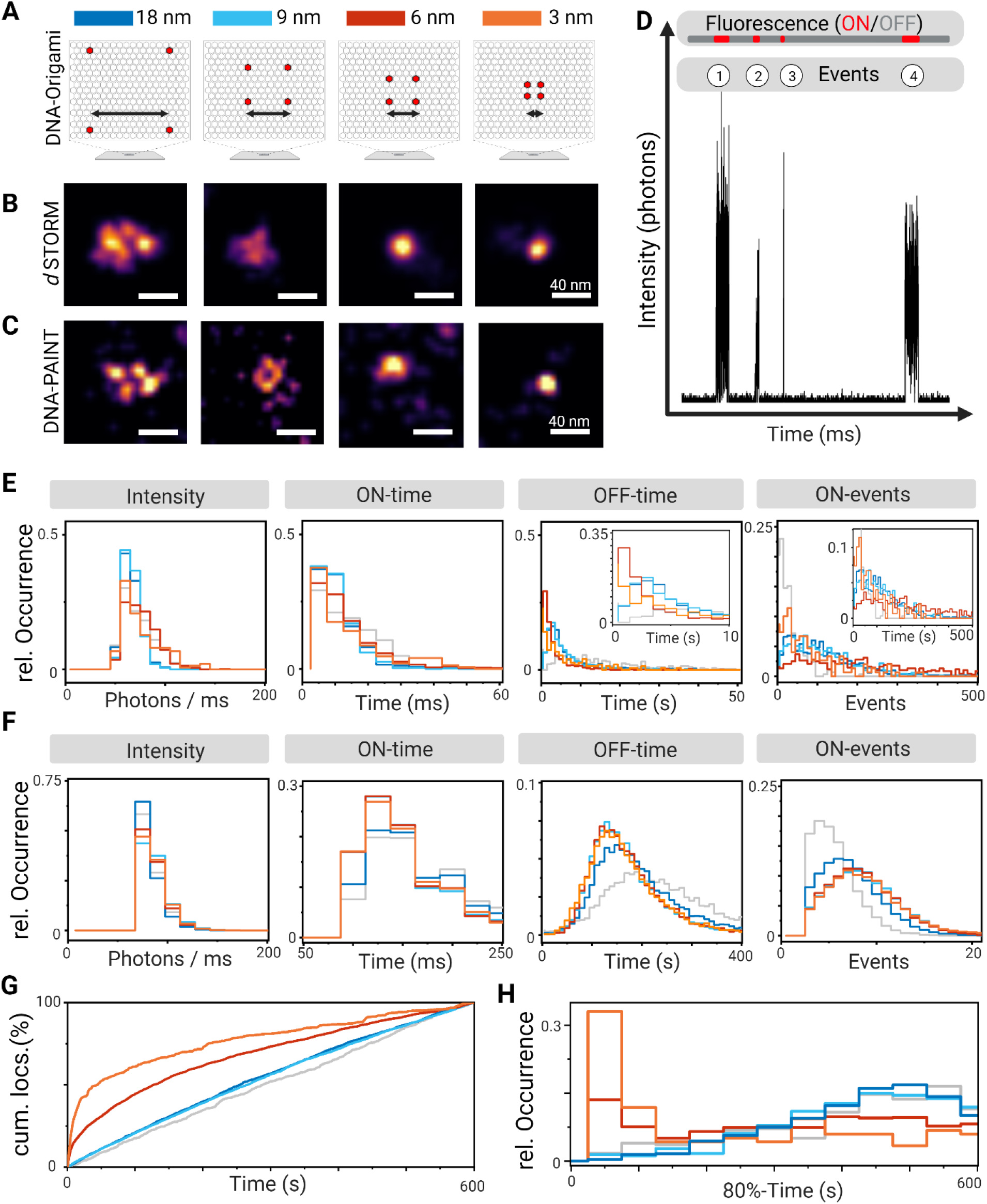
*d*STORM and DNA-PAINT imaging of DNA origami. (**A**) Scheme of DNA origami labeled with four Cy5 at interfluorophore distances of 18, 9, 6, and 3 nm. (**B** and **C**) Selected *d*STORM and DNA-PAINT images of DNA origami. (**D**) Analysis of fluorescence trajectories recorded from individual DNA origami imaged using 640 nm excitation at an intensity of 5 kW cm^-2^. (**E** and **F**) Relative occurrence of fluorescence intensity ms^-1^ in the on-state (Intensity), lifetime of the on-state (On-time), lifetime of the off-state (Off-time), and number of on-states (On-events) detected for DNA origami with different interfluorophore distances in (E) *d*STORM and (F) DNA-PAINT experiments (Color code: singly labeled reference (grey), 18 nm (dark blue), 9 nm (light blue), 6 nm (red), 3 nm (orange)). (**G**) Number of on-events (cumulative localizations, cum. locs.) detected per frame as a function of time during 10 min *d*STORM movies (movies S6-S11) of DNA origami with different interfluorophore distance. (**H**) Histogram of the times after which 80% of all localizations were detected per individual DNA origami.

In order to understand what causes the changes in photoswitching kinetics we have to revisit the *d*STORM switching mechanism. *d*STORM temporally separates the fluorescence of individual organic dyes by transferring the majority of them into a nonfluorescent off-state at the beginning of the experiment upon irradiation with intensities of a few kW cm^-2^ in thiol-based photoswitching buffer (*6,7*). The fluorescent on-state of a small subset of fluorophores is then generated by irradiating the sample usually at shorter wavelengths, i.e. typically at 405 nm. Unfortunately, there is no real consensus as to the origin of photoswitching of the two favorite *d*STORM cyanine dyes AF647 and Cy5 (*19,20*). While a recent study identified the formation of a Cy5-thiol adduct with absorption maximum at 310 nm as off-state in *d*STORM experiments (*21*)another study proposed the formation of a radical formed by one-electron reduction of the cyanine dye with a lifetime of a few tens of milliseconds and an absorption maximum at ~450 nm (*22*). But, as a matter of fact, Cy5 fluorescence can be restored from the off-state also upon irradiation with red light demonstrating that the off-state exhibits a very broad absorption spectrum (*21*). This also corroborates the experimental finding that *d*STORM imaging can be performed using exclusively irradiation at the absorption maxima of AF647 and Cy5 (*2,11*).

These considerations indicate that the nonfluorescent off-state of fluorophores can serve as energy transfer acceptor for the on-state of other fluorophores, i.e. for interfluorophore distances <10 nm the on-state of one fluorophore can serve as donor and excite fluorophores residing in their off-state (acceptors) via fluorescence resonance energy transfer (FRET) (*23,24*) into higher excited states from which the on-state can be repopulated. Because FRET from a donor with emission maximum at ~670 nm to an acceptor with a low extinction coefficient in the red wavelength range is inefficient, it is difficult to detect by standard means, e.g. fluorescence quenching of the donor (on-state). However, with increasing number of off-states present in the near-field of a donor the impact of these energy transfer processes on the fluorescence behavior of the multichromophoric system increases and will be measurable. In addition, albeit inefficient, each successful FRET event that transfers a fluorophore from the off-to the on-state will change the blinking pattern of the multichromophoric systems. Therefore, we hypothesized that photoswitching kinetics should directly report about the interfluorophore distance in the sub-10 nm range. In practice, this means that photoswitching should be accelerated at shorter interfluorophore distance and result in accumulation of fluorophores in their on-state.

Multichromophoric systems composed of several fluorophores separated by less than 10 nm show very complex fluorescence trajectories including collective off-states and different intensity levels also in the absence of photoswitching buffer because the fluorophores can interact by various energy transfer pathways including energy hopping, singlet-singlet- and singlet-triplet-annihilation. Therefore, multichromophoric systems often behave like single emitters in photon antibunching experiments (*25–27*). In addition, red-absorbing cyanine dyes such as Cy5 show a peculiar complicated behavior because of photoinduced isomerization from the fluorescent *trans* to a nonfluorescent *cis* state and back-isomerization (*28*). In addition, both the absorption spectra of the triplet state (*λ_max_* = 695 nm, *ε* = 105.000 cm^-1^ M^-1^) and *cis* state (*λ_max_* = 675 nm, *ε* = 326.000 cm^-1^ M^-1^) overlap strongly with the fluorescence emission of Cy5 (*21*). Considering the fact that Cy5 spends ~50% of the time in its *cis* state under equilibrium conditions in aqueous solutions that exhibits a lifetime of ~200 μs (*21,28*)it becomes obvious that Cy5 fluorophores separated by less than 10 nm can interact by various energy transfer pathways that result in the observation of blinking processes on different time scales (*21, 25–27*).

### Consequences for sub-10 nm fluorescence imaging

To elucidate how the described energy transfer processes compromise super-resolution microscopy in the sub-10 nm range we analyzed the fluorescence trajectories recorded in *d*STORM experiments from individual DNA origami with different interfluorophore distance at a temporal resolution of 5 ms (Fig. 1D). We were especially interested to find out if the blinking pattern of individual fluorophores, termed in the following photoswitching fingerprints, can be used to decode information about the underlying multichromophoric system. Analysis of individual photoswitching fingerprints revealed that the on-state of multiple labeled DNA origami shows similar lifetimes (on-time) but shorter off-state lifetimes (off-times) compared to the single dye reference as expected for multiple blinking fluorophores (Fig. 1E). In addition, the data clearly point out that the off-times decrease with decreasing interfluorophore distance (see enlarged section in Fig. 1E), i.e. the photoactivation rate increases with decreasing interfluorophore distance as expected for energy transfer between the on- and off-state. Furthermore, the on-state intensity is identical for the reference and the 18 and 9 nm DNA origami but slightly higher for the shorter interfluorophore distances (Fig. 1E). In contrast, DNA-PAINT experiments revealed photoswitching parameters independent of the distance of docking strands. Only the singly labeled reference shows, as expected longer off-times and less on-events (Fig. 1F). This result is expected since the binding of imager to docking strands is sequential in time and decoupled from irradiation. The broad distributions of the number of on-events indicates that for the majority of DNA origami less than four fluorophores and docking strands, respectively, are localized (Figs. 1E and 1F).

The low localization probability in both, *d*STORM and DNA-PAINT experiments might be explained by incomplete incorporation of modified oligonucleotides and labeling, respectively. In addition, in DNA-PAINT steric hindrance of docking and imager strands with lengths of 11 and 10 bases can distort labeling and transient binding especially at shorter distances. In *d*STORM experiments fast blinking observed as flickering at the very beginning of irradiation (movies S9-S11) promotes fast photobleaching and might thus impede the localization of all fluorophores as individual emitters.

Another way of looking at the blinking statistics of DNA origami is to plot the summed up localizations detected per frame as a function of time. Here we see that the singly labeled reference and the 9 and 18 nm DNA origami show a linear increase in the number of localizations with time in *d*STORM experiments (Fig. 1G). In a DNA origami carrying a single or four non-communicating Cy5 dyes each fluorophore will reside on average for several milliseconds in the on-state and several seconds in the off-state. Accordingly, homogeneous blinking is observed until photobleaching occurs. At shorter distances, however, the DNA origami show substantially faster blinking due to energy transfer from the on- to the off-state and subsequent repopulation of the on-state. The distribution of times after which 80% of all localizations are detected per DNA origami confirms fast blinking during the first minutes for the majority of the 3 and 6 nm DNA origami (Fig. 1H and fig. S5). Thus, our data clearly show that the temporal development of localizations detected from a sample labeled with Cy5 fluorophores changes at interfluorophore distances of <10 nm.

The consequences of our findings for sub-10 nm fluorescence imaging are apparent considering that at the very beginning of a *d*STORM experiment all fluorophores reside in their fluorescent on-state and have to be transferred to their off-state upon irradiation. During this time fast photoswitching can be initiated and fluorophores might be photobleached already during the first few tens of seconds of the experiment, e.g. during sample alignment. Consequently this results in substantially decreased localization probabilities and lower structural resolutions just like as it has been observed in previous structured illumination single-molecule localization experiments (*8–12*).

### Time-resolved fluorescence detection reveals the average interfluorophore distances

To obtain a more detailed picture of the photoswitching characteristics, we investigated individual DNA origami with higher temporal resolution by time-resolved confocal single-molecule fluorescence microscopy (Figs. 2A-E). Single-molecule surfaces were scanned with very low irradiation intensity to minimize premature photobleaching, individual DNA origami selected and parked in the laser focus to record the fluorescence intensity and lifetime with time. In photoswitching buffer, the reference DNA origami displayed blinking with short (milliseconds) on-states and long (seconds) off-states (Fig. 2A and fig S6). As expected for four independently emitting fluorophores 18 nm DNA origami displayed more switching events per time with similar intensities (Fig. 2B and fig. S7). The 9 nm DNA origami showed very similar behavior, however, some parts of the trajectories indicated faster photoswitching kinetics especially at the beginning of irradiation (Fig. 2C and fig. S8). In contrast, some of the 6 and 3 nm DNA origami showed clearly very fast photoswitching kinetics at the very beginning of the trajectories. Magnified views of the first seconds of the trajectories emphasize that the off-state lifetimes of DNA origami with interfluorophore distances of 3 and 6 nm are much shorter, i.e. blinking occurs on the millisecond time scale (Figs. 2D, 2E and figs. S9, S10).

**Fig. 2.**
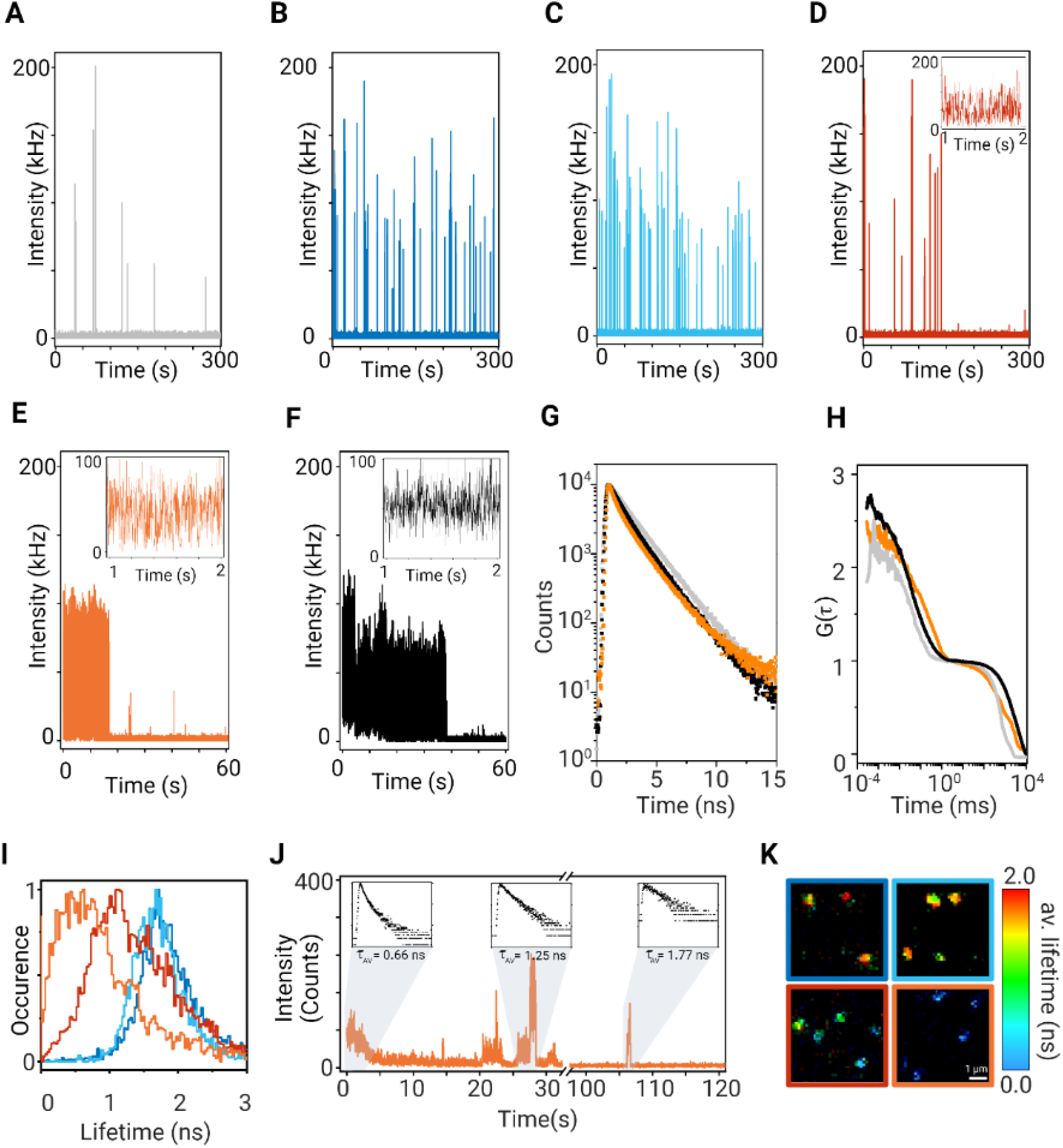
Various energy transfer pathways are responsible for fast blinking observed in the sub-10 nm range. (**A-E**) Fluorescence trajectories recorded for singly-labeled reference (A), 18 nm (B), 9 nm (C), 6 nm (D), and 3 nm (E) DNA origamis in *d*STORM photoswitching buffer. (Color code: singly labeled reference (grey), 18 nm (dark blue), 9 nm (light blue), 6 nm (red), 3 nm (orange)). Zoomed-in trajectories of the first seconds show fast blinking observed for the 6 and 3 nm DNA origamis. Time bins, 1 ms. (**F**) Fluorescence trajectory recorded for a 3 nm DNA origami in trolox buffer and zoomed-in fluorescence signals of the first two seconds. Time bin, 1 ms. (**G**) Average fluorescence decays form n=7-10 individual fluorescence trajectories of a singly labeled reference and the 3 nm DNA origamis measured in trolox and photoswitching buffer, respectively, revealing different energy transfer pathways between the Cy5 fluorophores. (**H**) Average intensity autocorrelation functions calculated from n=7-10 individual fluorescence trajectories of a singly labeled reference and the 3 nm DNA origamis measured in trolox and photoswitching buffer, respectively, normalized to 1 ms. (**I**) Histogram of average fluorescence lifetimes measured for individual DNA origami with different interfluorophore distances of 18, 9, 6, and 3 nm in photoswitching buffer. (**J**) Fluorescence trajectory of a 3 nm DNA origami in photoswitching and corresponding fluorescence decays with average fluorescence lifetimes of 0.66 ns, 1.25 ns, and 1.77 ns recorded during the gray marked areas. (**K**) Fluorescence lifetime (FLIM) images of the 18, 9, 6, and 3 nm DNA origami measured in trolox buffer emphasize the increased blinking and shorter fluorescence lifetime of Cy5 fluorophores in the sub-10 nm range (moving from top left to the bottom right). The samples were excited at 640 nm with 2.5 kW cm^-2^ at an integration time of 5 μs pixel^-1^).

In addition, we performed photon antibunching experiments to investigate the number of emitting fluorophores contributing to the detected fluorescence signal per single DNA origami (figs. S6-S10). Photon antibunching experiments take advantage of the fact that the probability of emitting two consecutive photons drops to zero for a single emitter for time intervals shorter than the excited-state lifetime. For sufficiently short laser pulses the number of photon pairs detected per laser pulse can be used to determine whether the emission is from one or more independently emitting quantum systems (*25–27,29*). Since the intensity of the central peak contains information about the number of independently emitting molecules, the number of photon pairs detected in the central peak, *N_c_*, at delay time zero, to the average number of the lateral peaks, *N_l,av_*, can be used to determine the number of independently emitting fluorophores. For example, neglecting background, *N_c_/N_l,av_* ratios of 0.0, 0.5, 0.67, and 0.75 are expected for 1-4 independently emitting fluorophores, respectively (*25–27,29*). If, for interfluorophore distances of < 10 nm the on-states are repopulated via energy transfer we expect a higher *N_c_/N_l,av_* ratio measured for the 3 and 6 nm DNA origami. And in fact, the *N_c_/N_l,av_* ratios increase from 0.067 (reference) via 0.073 (18 nm) and 0.085 (9 nm) to 0.207 (6 nm) and 0.255 (3 nm) for the different DNA origami (fig. S11). Higher *N_c_/N_l,av_* ratios are prevented by efficient energy hopping, singlet-singlet- and singlet-triplet-annihilation between Cy5 fluorophores in the on-state (*25–27*). Nonetheless, the slightly increased ratios detected for sub-10 nm interfluorophore distance are in accordance with the slightly higher fluorescence intensities recorded in *d*STORM experiments form the 3 and 6 nm DNA origami (Fig. 1E).

To dissect the two different energy transfer pathways (*trans*/*cis* and on/off) we investigated the 3 nm DNA origami in the absence of photoswitching, i.e. in PBS, pH 7.6 containing 1 mM trolox/troloxquinone and an oxygen scavenging system to prolong the observation time (*30*). The fluorescence trajectories of the 3 nm DNA origami showed similar blinking behavior in trolox buffer during the first seconds of irradiation but more fluorescence intensity levels (Fig. 2F and fig. S12). The similarity of the trajectories in the absence and presence of photoswitching buffer corroborates our hypothesis that in *d*STORM experiments with interfluorophore distances of <10 nm all fluorophores are efficiently transferred to the on-state resulting in fast blinking or flickering dependent on the temporal resolution of the experiment. Average fluorescence decays generated from n=7-10 fluorescence trajectories of individual reference and 3 nm DNA origami recorded in aqueous buffer by time-correlated single-photon counting (TCSPC) (figs. S6, S10, S12) clearly revealed a fluorescence quenching pathway at 3 nm interfluorophore distance indicated by a short fluorescence lifetime component (Fig. 2G). Contrary, the reference DNA origami labeled with a single Cy5 displayed a monoexponential fluorescence lifetime of ~1.8 ns. The fluorescence decay of the 3 nm DNA origami recorded in trolox buffer exhibited multiexponential kinetics with a short lifetime component of ~600 ps due to energy transfer from one fluorophore in the *trans* to another fluorophore residing in the *cis* state, neglecting energy transfer to the shorter-lived triplet state (Fig. 2G). As a photoinduced process, the efficiency of *trans/cis* isomerization, i.e. the degree of energy transfer, is determined by the irradiation intensity and thus not seen in standard ensemble TCSPC experiments where low irradiation intensities are usually applied (fig. S13).

Intriguingly, the fluorescence decay of 3 nm DNA origami recorded in photoswitching buffer exhibited multiexponential kinetics with a shorter fluorescence lifetime component of ~400 ps (Fig. 2G). The shorter fluorescence lifetime component confirms the additional energy transfer pathway in photoswitching buffer from one fluorophore in the on-state (donor) to another fluorophore in the off-state (acceptor). Furthermore, direct comparison of the fluorescence intensity autocorrelation functions of 3 nm DNA origami recorded during the first seconds of the trajectories demonstrates that the fluorescence fluctuations are dominated by energy transfer from the fluorescent *trans* to the nonfluorescent triplet and the *cis* state of Cy5 in trolox buffer. However, in photoswitching buffer an additional small on/off component appears in the few hundred microseconds range which we attribute to energy transfer from the on- to the off-state followed by repopulation of the on-state (Fig. 2H). Since all the observed on/off processes are strongly controlled by the excitation efficiency, reduction of the irradiation intensity slows down the blinking kinetics but simultaneously decreases the fluorescence intensity in the on-state and thus the localization precision (fig. S14).

In conclusion, the interfluorophore distance determines the off-state lifetime in *d*STORM experiments and is also encoded in the fluorescence lifetimes, whereas the number of on-events detected contains information about the number of fluorophores present. Consequently, the fluorescence lifetime of DNA origamis decreases with decreasing interfluorophore distance (Fig. 2I). Furthermore, the fluorescence lifetime of fluorophores in the 3 nm DNA origami increases during the fluorescence trajectory with progressing fluorophore photobleaching (Fig. 2J). In the example shown, the average fluorescence lifetime increases from 0.66 ns at the very beginning via 1.25 ns to 1.77 ns at the end when all fluorophores but a single survived (Fig. 2J and fig. S15). This demonstrates that the quenching efficiency of the on-state is determined by the number of off-states (quenchers) present. Fluorescence lifetime (FLIM) images of the four DNA origami measured in trolox buffer confirm that fast photobleaching of fluorophores during the first seconds of irradiation impede the observation of the described energy transfer processes. While individual 18 and 9 nm DNA origami are imaged with lifetimes of ~2 ns, 6 and 3 nm DNA origami show partially patchy spots with substantially shorter fluorescence lifetimes (Fig. 2K and fig. S16).

### Sub-10 nm super-resolution fluorescence imaging in cells

To translate our findings into biological applications, i.e. super-resolution imaging in cells, the labeling problem has to be solved first. While site-specific and efficient labeling of DNA origami with organic dyes is straightforward, site-specific fluorescence labeling of biomolecules separated by only a few nanometers remains challenging. In addition, the displacement of the fluorophore from the point of interest (the linkage error) and the conformational flexibility of the linker determine the localization accuracy achievable in super-resolution imaging experiments. Approaches to minimize the displacement of the fluorophore have been introduced including nanobodies and peptide tags but still yield linkage errors of a few nanometers (*31,32*) thus preventing the translation of 1-5 nm localization precision into image resolution in real biological samples. Furthermore, the sheer size of the fluorescent probe including fluorophore, linker, and affinity reagent does not only increase the linkage error but also limits the achievable labeling density (*33,34*). One approach to solve the labeling problem is direct covalent site-specific attachment of an organic dye to a protein of interest, which can be achieved by genetic code expansion (GCE) incorporating a non-canonical amino acids (ncAAs) into the protein of interest that can be efficiently labeled by bioorthogonal click chemistry with small organic dyes (*35,36*). The method enables site-specific efficient labeling of intra- and extracellular proteins with a linkage error of ~1 nm with super-resolution microscopy suited organic dyes (*37*). Latest studies conclusively demonstrated that click labeling of non-natural amino acids with small tetrazine-dyes is a versatile tool for the labeling of sterically difficult to access protein sites also in crowded environments (*38,39*).

We hypothesized that the combination of time-resolved photoswitching fingerprint analysis in combination with GCE with ncAAs and click-labeling can be used to unravel information about the molecular stoichiometry and interfluorophore distances in the sub-10 nm range in biological samples. We selected two different multimeric proteins, the hetero-pentameric γ-aminobutyric acid type A (GABA-A) (*40*) and the tetrameric kainate receptor (GluK2) (*41*). Site-specific labeling was achieved by incorporation of one or more trans-cyclooct-2-ene (TCO)-modified ncAAs (TCO*-L-lysine) into the extracellular domains of the (*i*) monomeric γ2-subunit, (*ii*) dimeric α2-subunit of GABA-A, and ( *iii*) homotetrameric GluK2 (Figs. 3A–3C). To identify the best positions for the insertion of ncAAs, various click constructs were generated: L198TAG and S217TAG for GABA-A receptors γ2-subunit, K73TAG, S171TAG, S173TAG, S181TAG, S201TAG, and K274TAG for GABA-A α2-subunit, and in addition to the previously described constructs S47TAG, S272TAG, S309TAG, S343TAG (*37*), we tested positions S398TAG, K494TAG and S741TAG for tetrameric GluK2 due to their rectangular positioning. All positions were selected to be at unstructured, extracellular regions of the pentameric GABA-A receptor (PDB-ID 6HUG) or homotetrameric GluK2 receptor (PDB-ID 5KUF). The generated click mutants were tested for ncAA incorporation and labeling efficiency in HEK293T cells (fig. S17).

**Fig. 3.**
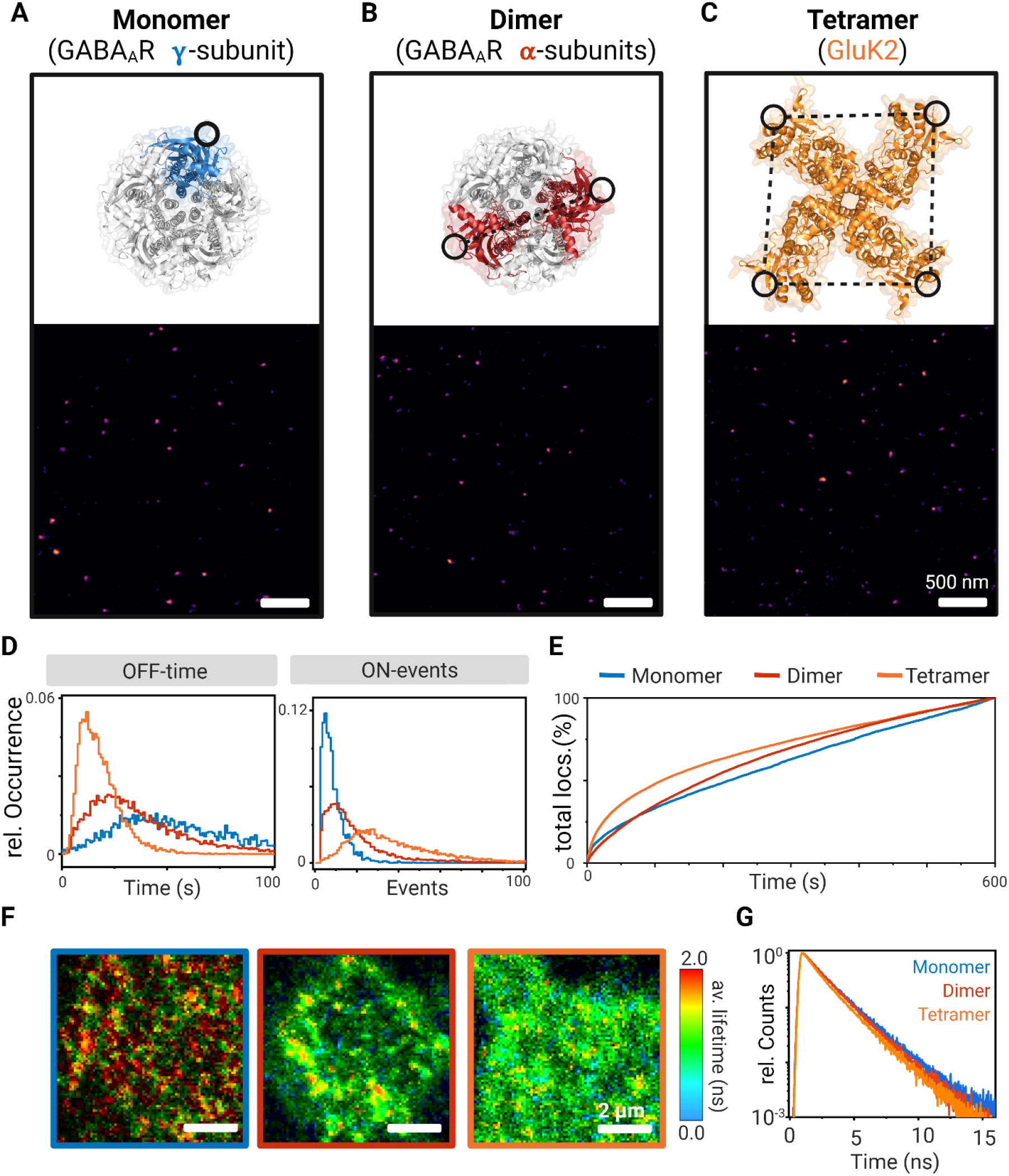
Time-resolved photoswitching fingerprint analysis in cells. (**A-C**) Molecular structures of the pentameric GABA-A (PDB-ID 6HUG) and tetrameric GluK2 receptor (PDB-ID 5KUF) with incorporation sites of ncAAs shown as black circles (blue: γ2-subunit GABA-A, red: dimeric α2 GABA-A, red: homotetrameric GluK2) and corresponding *d*STORM images of HEK293T membrane sections shown fluorescence signals of individual receptors (5 nm pixel^-1^). The ncAAs were labeled by click chemistry with Met-Tet-Cy5. In the GABA-A^S181TAG^ mutant the distance between the two fluorophores in the α2-subunits is ~5 nm. In the GluK2^S398TAG^ mutant the distance between the four Cy5 molecules is ~7 nm (*41*). (**D**) Relative occurrence of lifetimes of the off-state (Off-time), and number of on-states (On-events) detected from individual receptors in *d*STORM experiments. (**E**) Number of on-events (localizations) detected per frame as a function of time during 10 min *d*STORM experiments of membrane receptors. (**F**) FLIM images of HEK293T cells expressing monomeric γ2-subunit of GABA-A (left), dimeric α2-subunit of GABA-A (middle), and homotetrameric GluK2 receptors click-labeled with Met-Tet-Cy5 measured by confocal TCSPC imaging in photoswitching buffer at an irradiation intensity of 2.5 kW cm^-2^. To minimize photobleaching of fluorophores FLIM images were recorded at 5 μs integration time per pixel. No intensity threshold was applied. (**G**) Average fluorescence decays form n=10 FLIM images of HEK293T cells expressing receptors labeled with one, two, and four Cy5 fluorophores.

TCO*-L-lysine (TCO*A) reacts with tetrazine-dyes in an ultrafast, specific, and bioorthogonal inverse electron-demand Diels-Alder reaction and allows thus efficient site-specific labeling of receptors with one, two, and four Me-Tet-Cy5 dyes, respectively, with minimal linkage error (*36–38*). While the distances between the two fluorophores in the α2-subunits of GABA-A^S181TAG^ is ~5 nm, the interfluorophore distances in the tetrameric GluK2^S398TAG^ is ~7 nm (Figs. 3A–3C and fig. S17) (*42*).Hence, *d*STORM cannot resolve the different fluorophores and the resulting images display homogenous distributions of the receptors with no indications of clustering or different labeling stoichiometry (Figs. 3A-C and fig. S18). Photoswitching fingerprint analysis of the receptor signals recorded in *d*STORM experiments shows that the number of on-events and the lifetime of the off-states are unequivocally different reflecting the different number of fluorophores present per membrane receptor. We detected on average ~7 and ~15 on-events (median) for singly (γ2) and double (α2) labeled GABA-A receptors, receptively, and ~33 on-events (median) for the fourfold labeled GluK2 receptor (Fig. 3D). Hence, the number of on-events contains information about the number of fluorophores present per spatially unresolvable area (e.g. the receptor stoichiometry) even though the fluorophores are separated by less than 10 nm. This result reports most impressively that site-specific TCO*A incorporation into subunits of multimeric proteins by GCE followed by bioorthogonal click labeling with tetrazine-dyes enables quantitative labeling of protein sites separated by only a few nanometers.

The temporal evolution of localizations (on-events) detected per frame displays that energy transfer between the on- and off-states of the four Cy5 fluorophores in the GluK2 receptor results in shortening of the off-state lifetime and correspondingly more frequent blinking during the first minutes of irradiation (Fig. 3E) similar to the observation for the 6 and 3 nm DNA origami (Fig. 1G). For the single and double labeled GABA-A receptor we could not detect unequivocal differences in the temporal evolution of localizations (Fig. 3E) demonstrating that energy transfer from an on- to a single off-state is impossible to detect in SMLM data at an interfluorophore distance ~5 nm. In the presence of three possible acceptors (GluK2), however, energy transfer to the off-state and repopulation of the on-state can be easily identified (Fig. 3E). To explore if energy transfer between the on- and off-state can be detected we performed again time-resolved fluorescence experiments. FLIM images of HEK293T cells in photoswitching buffer clearly demonstrated that the singly labeled γ2 GABA-A receptor exhibits a monoexponential lifetime of ~1.85 ns independent of the irradiation intensity whereas GluK2 receptors exhibit a shorter fluorescence lifetime because of energy transfer from the *trans* on- to the off- and *cis* state (Fig. 3F and fig. S19). FLIM images of the double-labeled α2 GABA-A receptor are, in accordance with the photoswitching fingerprint analysis (Figs. 1D and 1E), less conclusive confirming that energy transfer from an on- to a single off-state remains difficult to be identified even by time-resolved fluorescence spectroscopy (Fig. 3F and fig. S19).

This impression is also supported by average fluorescence decays (fig. S19). Here, both the average fluorescence decays of α2 GABA-A and GluK2 exhibit multiexponential character with short fluorescence lifetime components of ~0.65 ns and ~0.75 ns and amplitudes of ~0.16 and ~0.38, respectively (Fig. 3G). This means that energy transfer between two Cy5 separated by ~5 nm at best induces the appearance of a small 650 ps component in the fluorescence decay. On the other hand, energy transfer between four Cy5 separated by ~7 nm causes a slightly longer lifetime component of ~750 ps but with higher amplitude and is thus far more easy to detect.

### Discussion and outlook

Since in *d*STORM experiments only a single fluorophore is expected to reside in the on-state per diffraction-limited area, being cycled between its singlet-ground and first excited singlet-state for several milliseconds, fluorophore interactions have been presumed to play a negligible role. Hence, a direct relation between fluorophore interactions and image resolution in SMLM experiments is not given. However, as we have shown here dipole-dipole induced energy transfer between the on-state and off-state of fluorophores accelerates photoactivation resulting in faster repopulation of the on-state, and in combination with other additional energy transfer pathways from the *trans* on-state of cyanine dyes to the *cis* and triplet state, in fast blinking. Such fast switching events are elusive in *d*STORM experiments because samples are usually irradiated at high intensity for a while to turn the majority of fluorophores into their off-state before data acquisition. Due to this premature photobleaching, the localization probability decreases substantially with decreasing interfluorophore distance in the sub-10 nm range. The energy transfer efficiency is controlled by the acceptor concentration, i.e. the number of fluorophores residing in the off-, *cis* and triplet state. Since these states are populated via the excited state of fluorophores, the observed energy transfer efficiency depends critically on the irradiation intensity (figs. S14 and S19). The effects described are unavoidable in *d*STORM experiments if sub-10 nm spatial resolutions are to be achieved because of the required high labeling density.

The influence of these energy transfer pathways on the achievable spatial resolution in the sub-10 nm range has thus up to now not been perceived by the superresolution fluorescence imaging community. Or to be precise, near-field fluorophore interactions that decrease the localization probability of fluorophores separated by less than 10 nm have not been considered so far. The resulting lower image quality sparked a debate about the potential of new structured illumination SMLM methods (*8–12*) for molecular resolution imaging (*13*) but the contradiction between high localization precision and low localization probability remained enigmatic. Our findings are also important for quantitative SMLM approaches. Since photoswitchable fluorophores can switch multiple times between the on- and off-state before they permanently photobleach repeated localizations from the same fluorophore are detected, which complicates quantifying numbers of molecules. Multiple methods have been developed to correct for blinking-caused artifacts to enable quantification of SMLM data. However, all approaches rely on interfluorophore distance independent photoswitching kinetics of fluorophores to group localizations that likely come from the same fluorophore (*43–45*). Thus, our results clearly identify a new limitation of quantitative SMLM at high labeling densities with interfluorophore distances in the sub-10 nm range.

On the other hand, one can take advantage of near-field interactions of fluorophores developing a completely new approach to reveal the number of fluorophores and their interfluorophore distance in the sub-10 nm range. The analysis is based on the finding that information about the number of fluorophores present and their interfluorophore distance is encoded in their photoswitching fingerprints. The number of on-events gives directly the number of fluorophores, provided the temporal resolution of the experiment is high enough to resolve fast blinking processes. Which will probably never be the case, since in the initial period where fast blinking occurs too many fluorophores are active to localize (spatially resolve) them individually. The distance can be derived from the off-state lifetime of the fluorophores (Figs. 1E and 3D), the temporal evolution of detected localizations (Figs. 1G and 3F), the photon antibunching signature (fig. S11) and the fluorescence lifetime of the fluorophores (Figs. 2G, 2I–2K, 3F, 3G and figs. S15, S16, S19). Since all these processes are strongly controlled by the excitation efficiency, reduction of the irradiation intensity slows down the blinking kinetics and enhances the probability to accurately analyze the photoswitching fingerprints but simultaneously decreases the fluorescence intensity in the on-state and thus the localization precision (fig. S14). Since rhodamine and oxazine dyes also enter off-states that absorb at shorter wavelengths and can be photoactivated (*20*), photoswitching fingerprint analysis must not remain limited to the red spectral range. The simpler photophysics, i.e. the absence of *trans/cis* isomerization can potentially simplify data analysis.

Our data show clearly that GCE and site-specific incorporation of ncAAs into proteins followed by click-labeling (*35–39*) volunteers as method of choice for biological superresolution microscopy in the sub-10 nm range because of the virtually quantitative labeling efficiency. Anticipating photowitching fingerprint analysis of endogenous proteins, a potential limitation of GCE with ncAAs is the overexpression of the protein of interest. However, new emergent genome editing tools such as CRISPR/Cas9 might enable site-specific incorporation of ncAAs into endogenous proteins. Furthermore, orthogonal ribosomes (*46,47*) in combination with quadruplet codons (*48*) will contribute significantly to reduce suppression of endogenous amber codons and improve GCE efficiency, and therefore enable quantitative insertion of multiple ncAA into the protein of interest. Our results also demonstrate that energy transfer between identical fluorophores (e.g. between the *trans* and *cis* state and on- and off-state) can be used to determine the interfluorophore distance. Hence two or more identical ncAAs can be incorporated at different sites into the same protein or multiprotein complex. Bioorthogonal click-labeling with the same fluorophore in combination with time-resolved single-molecule fluorescence spectroscopy can then be used advantageously for distance measurements akin to standard FRET investigations.

Finally, the disclosure of fast blinking as a result of energy transfer between on- and off-states of fluorophores provides guidance how to further improve sub-10 nm fluorescence imaging. For example, GCE and bioorthogonal click labeling in combination with confocal fluorescence lifetime *d*STORM (*49*) and photoswitching fingerprint analysis might evolve as a powerful superresolution microscopy method for imaging in the sub-10 nm range. But also the advancement of MINFLUX (*8,12*) and especially pulsed interleaved MINFLUX (*50*) will benefit from our findings. The only possibility to avoid the acceleration of photoswitching rates and accumulation of dyes in the on-state are the use of DNA-PAINT (*18*) where only one imager strand is present per 10 nm area simultaneously during the experiments, or fluorophores whose switching mechanism is independent of irradiation, e.g. spontaneously blinking dyes such as the Si-rhodamine dye HMSiR (*51*) or bridged Cy5B under reductive imaging conditions (*52*). Their use in refined SMLM methods (*8–12*) might thus represent the method of choice to improve the localization probability and thus enable reliable sub-10 nm fluorescence imaging. However, HMSiR exhibits pH-dependent blinking properties and lower localization precision (*37,51*). DNA-PAINT achieves a higher spatial resolution than *d*STORM (Fig. 1C and fig S4) but requires site-specific labeling with docking strands with a length of ~10 DNA bases. It remains to be tested how the labeling efficiency with negatively charged oligonucleotides that repel each other can be optimized to ensure quantitative localization of protein sites at distances of only a few nanometers. Another possibility to avoid energy transfer between adjacent fluorophores is to expand the sample before imaging (*53*). Post-labeling expansion microscopy combined with SMLM improves the labeling efficiency and reduces the linkage error thus paving the way for superresolution fluorescence imaging with true molecular resolution (*54*). Although there is ample scope for alternative improvements, the run for an efficient sub-10 nm fluorescence imaging method has just begun.

## Acknowledgments

The authors thank C. Stigloher from the Department of Electron Microscopy, Biocenter, University of Würzburg for quality control of DNA origami structures and S. Reinhard for technical assistance with data analysis. Figures were created with biorender.com. The plasmid for the expression of GluK2 was a kind gift from P. Seeburg. GABA(A) receptor subunit a2SE was a gift from Tija Jacob & Stephen Moss (Addgene plasmid #49169). The plasmid for expressing the modified γ2 subunit of the GABA-A receptor in mammalian cells was kindly provided by A. Barberis. The plasmid for the expression of the tRNA/aminoacyl transferase pair (pCMV tRNAPyl/NESPylRSAF, herein termed PylRS/tRNAPyl) was kindly provided by E. Lemke.

## Author contributions

D.A.H., G.B., S.D. and M.S. conceived and designed the project. M.S. supervised the project. D.A.H. performed all *d*STORM and confocal single-molecule experiments. M.M. performed the DNA-PAINT measurements. G.B. designed the GCE experiments. G.B., D.T., A.K., M.St., performed ncAAs incorporation and click-labeling. D.A.H. and S.D. performed data analysis. M.S. wrote the manuscript. All authors revised the final manuscript.

## Data and materials availability

All data that support the findings described in this study are available within the manuscript and the related supplementary information, and from the corresponding authors upon reasonable request.

## Supplementary Materials

Materials and Methods

Figures S1-S19

Movies S1-S11

References (55–67)

## References

1. L. Schermelleh, et al. Super-resolution microscopy demystified. Nat. Cell Biol. 21, 72–84 (2019).

2. M. Sauer, M. Heilemann, Single-molecule localization microscopy in eukaryotes. Chem. Rev. 117, 7478–7509 (2017).

3. H. Deschout, et al., Precisely and accurately localizing single emitters in fluorescence microscopy. Nat. Methods 11, 253–266 (2014).

4. K. I. Mortensen, L. S. Churchman, J. A. Spudich, H. Flyvberg, H. Optimized localization analysis for single-molecule tracking and super-resolution microscopy. Nat. Methods 7, 377–381 (2010).

5. U. Endesfelder, S. Malkusch, F. Fricke, M. Heilemann, M. A simple method to estimate the average localization precision of a single-molecule localization microscopy experiment. Histochem. Cell Biol. 141, 629–638 (2014).

6. M. Heilemann, et al. Subdiffraction-resolution fluorescence imaging with conventional fluorescent probes. Angew. Chem. Int. Ed. 47, 6172–6176 (2008).

7. S. van de Linde, A. Löschberger, T. Klein, M. Heidbreder, S. Wolter, M. Heilemann, M. Sauer, Direct stochastic optical reconstruction microscopy with standard fluorescent probes. Nat. Protocols 6, 991–1009 (2011).

8. F. Balzarotti, et al. Nanometer resolution imaging and tracking of fluorescent molecules with minimal photon fluxes. Science 355, 606–612 (2017).

9. L. Gu, et al. Molecular resolution imaging by repetitive optical selective exposure. Nat. Methods 16, 1114–1118 (2019).

10. L. Reymond, et al. SIMPLE: structured illumination based point localization estimator with enhanced precision. Opt. Express 27, 24578–24590 (2019).

11. J. Cnossen, et al. Localization microscopy at doubled precision with patterned illumination. Nat. Methods 17, 59–63 (2020).

12. K. C. Gwosch, J. K. Pape, F. Balzarotti, P. Hoess, J. Ellenberg, J. Ries, S. W. Hell, MINFLUX nanoscopy delivers 3D multicolor nanometer resolution in cells. Nat. Methods 17, 217–224 (2020).

13. K. Prakash, A. P. Curd, Assessment of 3D MINFLUX data for quantitative structural biology in cells. bioRxiv https://doi.org/10.1101/2021.08.10.455294.

14. J. Engelhardt, J. Keller, P. Hoyer, M. Reuss, T. Staudt, S. W. Hell, S. W. Molecular orientation affects localization accuracy in superresolution far-field fluorescence microscopy. Nano Lett. 11, 209–213 (2010).

15. M. P. Backlund, et al. Removing orientation-induced localization biases in single-molecule microscopy using a broadband metasurface mask. Nat. Photonics 10, 459–462 (2016).

16. P. W. Rothemund, Folding DNA to create nanoscale shapes and patterns. Nature 440, 297–302 (2006).

17. J. J. Schmied, A. Gietl, P. Holzmeister, C. Forthmann, C. Steinhauer, T. Dammeyer, P. Tinnefeld, Fluorescence and super-resolution standards based on DNA origami. Nat. Methods 9, 1133–1134 (2012).

18. R. Jungmann, C. Steinhauer, M. Scheible, A. Kuzyk, P. Tinnefeld, F. C. Simmel, Single-molecule kinetics and super-resolution microscopy by fluorescence imaging of transient binding on DNA origami. Nano Lett. 10, 4756–4761 (2010).

19. G. T. Dempsey, M. Bates, W. E. Kowtoniuk, D. R. Liu, R. Y. Tsien, X. Zhuang, X. Photoswitching mechanism of cyanine dyes. J. Am. Chem. Soc. 131, 18192–18193 (2009).

20. S. van de Linde, I. Krstic, T. Prisner, S. Doose, M. Heilemann, M. Sauer, Photoinduced formation of reversible dye radicals and their impact on super-resolution imaging. Photochem. Photobiol. Sci. 10, 499–506 (2011).

21. Y. Gidi, L. Payne, V. Glembockyte, M. S. Michie, M. J. Schnermann, G. Cosa, Unifying mechanism for thiol-induced photoswitching and photostability of cyanine dyes. J. Am. Chem. Soc. 142, 12681–12689 (2020).

22. Lisovskaya, A., Carmichael, I., Harriman, A. Pulse radiolysis investigation of radicals derived from water-soluble cyanine dyes: implications for super-resolution microscopy. J. Phys. Chem. A 125, 5779–5793 (2021).

23. T. Förster, Zwischenmolekulare Energiewanderung und Fluoreszenz. Annu. Phys. 2, 55–75 (1948).

24. G. D. Scholes, Long–range resonance energy transfer in molecular systems. Annu. Rev. Phys. Chem. 54, 57–87 (2003).

25. P. Tinnefeld, K. D. Weston, T. Vosch, M. Cotlet, J. Hofkens, F. C. Dr Schryver, M. Sauer, Antibunching in the emission of a tetrachromophoric dendritic system. J. Am. Chem. Soc. 124, 14310–14311 (2002).

26. J. Hofkens, et al. Revealing Competitive Förster-Type Resonance Energy Transfer Pathways in Single Bichromophoric Molecules. Proc. Natl. Acad. Sci. USA 100, 13146–13151 (2003).

27. P. Tinnefeld, D.-P. Herten, S. Masuo, T. Vosch, M. Cotlet, J. Hofkens, K. Müllen, F. C. De Schryver, M. Sauer, Higher excited state photophysical pathways in multichromophoric systems revealed by single-molecule fluorescence spectroscopy. ChemPhysChem 5, 1786–1790 (2004).

28. J. Widengren, P. Schwille, Characterization of photoinduced isomerization and back-isomerization of the cyanine dye Cy5 by fluorescence correlation spectroscopy. J. Phys. Chem. A 104, 6416–6428 (2000).

29. B. Lounis, W. E. Moerner, Single photons on demand from a single-molecule at room temperature. Nature 407, 491–493 (2000).

30. T. Cordes, J. Vogelsang, P. Tinnefeld, On the mechanism of trolox as antiblinking and antibleaching reagent. J. Am. Chem. Soc. 131, 5018–5020 (2009).

31. I. Chamma, et al. Mapping the dynamics and nanoscale organization of synaptic adhesion proteins using monomeric streptavidin. Nat. Commun. 7, 10773 (2016).

32. D. Virnat, et al. A peptide tag-specific nanobody enables high quality labeling for dSTORM imaging. Nat. Commun. 9, 930 (2018).

33. C. E. Shannon, Communication in the presence of noise. Proc. IEEE Inst. Electr. Electron Eng. 37, 10–21 (1949).

34. W. R. Legant, L. Shao, J. B. Grimm, T. A. Brown, D. E. Milkie, B. B. Avants, L. D. Lavis, E. Betzig, High-density three-dimensional localization microscopy across large volumes. Nat. Methods 13, 359–365 (2016).

35. J. A. Prescher, C. R. Bertozzi, Chemistry in living systems. Nat. Chem. Biol. 1, 13–21 (2005).

36. C. C. Liu, P. G. Schultz, Adding new chemistries to the genetic code. Annu. Rev. Biochem. 79, 413–444 (2010).

37. G. Beliu, et al. Bioorthogonal labeling with tetrazine-dyes for super-resolution microscopy. Commun. Biol. 2, 261 (2019).

38. G. Beliu, et al. Tethered agonist exposure in intact adhesion/class B2 GPCRs through intrinsic structural flexibility of the GAIN domain. Mol. Cell 81, 905–921 (2021).

39. D. Bessa-Neto, et al. Bioorthogonal labeling of transmembrane proteins with non-canonical amino acids allows access to masked epitopes in live neurons. Nat. Commun. 12, 6715.

40. W. Wisden, P. H. Seeburg, GABAA receptor channels: from subunits to functional entities. Curr. Opin. Neurobiol. 2, 263–269 (1992).

41. J. Lerma, Roles and rules of kainate receptors in synaptic transmission. Nat. Rev. Neuroscience 4, 481–495 (2003).

42. J. R. Meyerson, S. Chittori, A. Merk, P. Rao, T. H. Han, M. Serpe, M. L. Mayer, S. Subramaniam. Structural basis of kainite subtype glutamate receptor desensitization. Nature 537, 567–571 (2016).

43. F. Baumgart, A. M. Arnold, K. Leskovar, K. Staszek, M. Fölser, J. Weghuber, H. Stockinger, G. J. Schütz, Varying label density allows artifact-free analysis of membrane-protein nanoclusters. Nat. Methods 13, 661–664 (2016).

44. G. Hummer, F. Fricke, M. Heilemann, Model-independent counting of molecules in single-molecule localization microscopy. Mol. Biol. Cell 27, 3637–3644 (2016).

45. C. H. Bohrer, X. Yang, S. Thakur, X. Weng, B. Tenner, R. McQuillen, B. Ross, M. Wooten, X. Chen, J. Zhang, E. Roberts, M. Lakadamyali, J. A. Xiao, A pairwise distance distribution correction (DDC) algorithm to eliminate blinking-caused artifacts in SMLM. Nat. Methods 18, 669–677 (2021).

46. O. Rackham, J. W. Chin, A network of orthogonal ribosome mRNA pairs. Nat. Chem. Biol. 1, 159–166 (2005).

47. K Wang, H. Neumann, S. Y. Peak-Chew, J. W. Chin, Evolved orthogonal ribosomes enhance the efficiency of synthetic genetic code expansion. Nat. BiotechnoK 25, 770–777 (2007).

48. H. Neumann, K. Wang, L. Davis, M. Garcia-Alai, J. W. Chin, Encoding multiple unnatural amino acids via evolution of a quadruplet-decoding ribosome. Nature 464, 441–444 (2010).

49. J. C. Thiele, D. A. Helmerich, N. Oleksiievets, R. Tsukanov, E. Butkevich, M. Sauer, O. Nevskyi, J. Enderlein, Confocal fluorescence-lifetime single-molecule localization microscopy. ACS Nano 14, 14190–14200 (2020).

50. L. A. Masullo, F. Steiner, J. Zähringer, L. F. Lopez, J. Bohlen, L. Richter, F. Cole, P. Tinnefeld, F. D. Stefani, Pulsed interleaved MINFLUX. Nano Lett. 21, 840–846 (2021).

51. S. N. Uno, M. Kamiya, T. Yoshihara, K. Sugawara, K. Okabe, M. C. Tarhan, H. Fujita, T. Funatsu, Y. Okada, S. Tobity, Y. A. Urano, A spontaneously blinking fluorophore based on intramolecular spirocyclization for live-cell super-resolution imaging. Nat. Chem. 6, 681–689 (2014).

52. M. S. Michie, et al. Cyanine conformational restraint in the far-red range. J. Am. Chem. Soc. 139, 12406–12409 (2017).

53. F. Chen, P. W. Tillberg, E. S. Boyden, Expansion Microscopy. Science 347, 543–548 (2015).

54. F. U. Zwettler, S. Reinhardt, D. Gambarotto, T. D. M. Bell, V. Hamel, P. Guichard, M. Sauer, Molecular resolution imaging by post-labeling expansion singlemolecule localization microscopy (Ex-SMLM). Nat. Commun. 11, 3388 (2020).

