## Supplementary Information for "Sub-10 nm fluorescence imaging"

**This PDF file includes:**

Materials and Methods

Figs. S1 to S19

Captions for Movies S1 to S11

**Other Supplementary Materials for this manuscript include the following:**

Movies S1 to S11

### MATERIALS AND METHODS

**Design, hybridization and quality control of DNA–origami structures.** DNA-origami rectangle structures were designed with caDNAno 2.2.0 (*55,56*). Stability calculations of the origami designed were performed using CanDO (*57,58*). All dye/TCO modified staple strands were ordered at biomers.net GmbH, whereas all biotinylated strands were ordered at SigmaAldrich Inc. All unmodified staple strands were ordered at Merck KGaA. We used the phage M13mp18 derivat DNA type p7560 as scaffold DNA (tilibit nanosystems, M1-32). Hybridization was done by mixing 10 nM scaffold DNA with 15x surplus of unmodified staple strands and 30x surplus of modified staple strands in hybridisation buffer, consisting of 5 mM Tris(hydroxymethyl)aminomethane (TRIS) (Merck, 1.08382.2500), 5 mM sodium chloride (NaCl) (Sigma, S5880-1KG), 1 mM ethylene diamine tetraacetic acid (EDTA) (Sigma, E1644-250G) and 12 mM magnesium chloride (MgCl2) (AppliChem, A4425,0500) using a ThermoCycler (C1000 Thermal Cycler, BioRad) with a linear thermal gradient of -1 °C/min from 90 °C to 4 °C. For DNA – PAINT Origami, trans-cyclooctene modified staple strands were used. These origami structures were incubated with a 10 fold surplus of docking strand 5´-modified with methyl - tetrazine (5´-3´: TTA TAC ATC TA, biomers.net) per TCO-staple for 2 h at 4 °C after hybridization. The hybridized samples were purified by electrophoresis in a 1.5 % agarose gel (Sigma, A9539-500G) in 1x TBE buffer, consisting of 4.5 mM TRIS (Merck, 1.08382.2500), 4.5 mM boric acid (Merck, K1898765) und 10 mM EDTA (Sigma, E1644-250G) and 0.5x TBE with 12 mM $MgCl_2$ (AppliChem, A4425,0500) as running buffer. After melting the agarose with a microwave, the solution was cooled down to ~60 °C until adding 12 mM MgCl2 (AppliChem, A4425,0500). Afterwards the gel was poured immediately. A small amount (~ 10 µl) of a sample was picked as reference, which was mixed with 2 µl intercalating dye (Safe-Green™, Applied Biological Materials Inc., G108-G). A small amount of pure scaffold as well as pure staple strands were mixed in hybridization buffer and used as references. These solutions were also mixed with intercalating dye. The rest of the hybridized Origami samples were not mixed with intercalating dye. All samples were mixed with loading dye, consisting of 10 mM TRIS (Merck, 1.08382.2500), 60 % glycerol (v/v) (Merck, 1.37028.1000) und 0.03 % bromophenol blue (w/v) (Carl Roth, T116.1). Electrophoresis was done at 70 V, using a programmable DC voltage source (PowerPac™ Basic, BioRad), for ~ 2h in water/ice bath. The part of the gel including the references were cut across the length of the gel and the bands marked at an UV transilluminator (UST20M-8E, INTAS). Afterwards, the marked gel was combined with the not illuminated part of the gel, containing the DNA-origami structures not mixed with intercalating dye. Not illuminated DNA-origamis were cut out according to the high of the marked references. The extracted gel parts were divided by cutting several times and purified via Freeze N´ Squeeze columns (Freeze N´ Squeeze, 7326165, BioRad) according the manufacturer instructions using a benchtop centrifuge (Biofuge fresco, Heraeus) at 13.000 g. For all measurements, the DNA-origami were produced freshly on the same day of the measurements. The shape and the quality of the purified DNA-origami structures were checked via transmission electron microscopy (JEM 1011, JEOL) and negative staining of the samples. Therefore carbon coated 100 Mesh TEM-grids were used and glowed freshly. The prepared grids were incubated with 15 µl sample solution for 2 minutes. Afterwards the solution was peeled of using a filter paper. The grid was dipped into a 0.75 % uranyl acetate solution (EMS, 22400) and peeled of immediately. This step was repeated 4 times until the grid was incubated with 0.75 % uranyl acetate solution (EMS, 22400) for 45 seconds. The solution was peeled of and air-dried.

**DNA origami sequences.** All dye modified staple sequences were also ordered unmodified

| Start | End | Sequence (5´ - 3´) | modification |
|---|---|---|---|
| 10[7] | 8[16] | CGAATTCGCCGGGTACCGATAGCATGTCAATCTACCTCGA | |
| 1[56] | 3[79] | GTGGATGTTCTTCTAAGTGGTTGTATATCCCATAATCGGC | |
| 9[184] | 11[175] | TATAACTACTTAGGTTGGGCACAAGAATTGAGAGAGACTA | |
| 17[184] | 19[175] | GTGAGTGAATAAATCAATAGAAACGTCACCAATACCTTTT | |
| 14[71] | 18[74] | ATAAATCAAGTACCTTTAATTGCTTTCGGT | 3´ - TCO/Cy5 (9nm) |
| 27[8] | 27[7] | ATGGTGGTAGAATAGCCCGAGATACCTGTTTG | |
| 23[48] | 20[56] | AACTGGCTCATTATACAATCAGGT | |
| 22[71] | 25[74] | CATAGTAAAGTATTAAGAG | 3´ - TCO/Cy5 (9nm) |
| 2[159] | 1[151] | GACGACAATAAACAACGAGCCAGT | |
| 21[80] | 16[88] | AATACTGCAAACGAGAATCACCGGAACCAGAGTATAACAG | |
| 31[16] | 28[24] | ATTAAAGAACGTGGACTCCCTTAT | |
| 25[80] | 20[88] | GACTCCTCACAGTTAAAAGAAAAATCTACGTTAGTTCAGA | |
| 26[167] | 29[183] | GGAGGTTTCGTAACGATCTAAAGTTTTGTTAA | |
| 13[24] | 10[8] | GTACCAAAGATGAACGGTAATCGTAAAACGCT | |
| 22[63] | 20[48] | GAGCAACACTATCATATAATAGTACTTTACCC | |
| 8[175] | 11[167] | CAAATCCAATCGCAAGTAGGTCTG | |
| 25[75] | 27[79] | GCTGACATTACCCGCTGGCTG | |
| 6[167] | 9[183] | TATTCTAAGCTAATATCAGAGAGATAACCTTA | |
| 25[88] | 22[80] | AAGAGAAGATAACGCCAAAAGGAA | |
| 25[120] | 21[127] | CAGTACCATTAGGAATACCACATTATCTGACAGGAGGTTG | |
| 13[88] | 10[84] | CAGTATGTTTTTTGAGAGAT | 3´ - TCO/Cy5 (6nm) |
| 30[47] | 32[24] | AGACTTTTGGCTACAGCAGCATCGGAACGAGGTCCAACGT | |
| 19[112] | 16[120] | CTCCCTCAGAGCCGCCACTAAAGT | |
| 28[183] | 28[184] | TTCAGGTTTTTACATCGGGAGAAACGTAGATT | |
| 21[184] | 23[175] | AAACATCAGAAGATGATGAAGCCAGAATGGAATTCATTTC | |
| 25[24] | 22[8] | TTGGGCTTCGACGATAAAAACCAAAATAGACC | |
| 30[39] | 32[48] | TCATGAGGAATTCGACAACTCGTATTAAATCCGCGAAAGA | |
| 5[88] | 2[80] | GTGCATCTCTGAACAAGAAAAATA | |
| 31[80] | 28[88] | CCAAAAGGAGCCTTTACATGTTAC | |
| 24[175] | 27[167] | ACAAAATCGCGCAGAGTACAGTAA | 5´ biotin |
| 26[143] | 31[135] | GAACCGCCCGTAACACTGTAGCATTCATCGCCGAATTTCT | |
| 14[7] | 12[16] | AAGTGTAAGAGCCGGAAGCAGCTAAATCGGTTCGCTCACA | |
| 17[88] | 14[80] | TAAGAGGTTAATAGTAGTAGCATT | |
| 14[111] | 17[101] | GGAAGGTAAATATTACCATTTT | |
| 21[24] | 19[15] | CAAAAGAATCAAATATCGCGTTTTAATTCCCTTAATGAAT | |
| 33[56] | 28[48] | TTTGCCCGTTTTCACGAGGACTAAGAAAGAGGAAGGGAAC | |
| 14[159] | 12[144] | TTTACCAGCGCCAAAGACGCAAAGAAGCCCTT | |
| 29[56] | 25[63] | CAAAAGAAGTAACAAAGCTGCTCACTCTGAAACATGAA | |
| 13[184] | 15[175] | AAGACGCTCGATAGCTTAGATCAATAGAAAATCCCTTAGA | |
| 20[175] | 23[167] | TACATTTAACAATTTCGCGAATTA | |
| 29[152] | 24[144] | ATAGTTAGAGTACCGCCACCCTCACGAGAGGGTACAGGAG | |
| 17[24] | 15[15] | AACCAGACCAATAAAGCCTCAGAGCATAAATAGTGCCTAA | |
| 3[144] | 0[152] | AGCCGTTTTTATTTTCTTCTTACC | |
| 5[24] | 3[15] | GAACAAACAGGGTTTTCCCAGTCACGACGTCTTGGGCACG | |
| 33[0] | 32[16] | TTGAGGATTTAGAAGTATTAGACTCAAAGGGC | |
| 6[83] | 7[79] | GCACCCAGCTACGCGTCTTT | |
| 23[16] | 20[24] | CCCTTCACCGCCTGGCAGAGGCGG | |
| 9[102] | 10[102] | AAGCATTAGACCGGAGAGGG | 3´ - TCO/Cy5 (6nm) |
| 11[16] | 6[16] | TGTTTCCTGTGTGAAAGTGCTTGTATATGTACGTGAGCGAGTAACAAC | |
| 29[88] | 26[80] | CCCCAGCGACAAGAACCGGATATT | |
| 5[56] | 2[48] | GTCACGTTTACGAGCAAAGGCGAT | |
| 15[48] | 12[53] | GTTTAGCTATATTTTCGAAGAGAAGCC | |

| | | | |
|---|---|---|---|
| 9[56] | 6[48] | CAAAAACACCTGAATCAGCCAGCT | 3´ - Cy5 (18nm) |
| 2[135] | 5[143] | CTAATGCATCAGGAAGATCGCACT | |
| 29[24] | 26[8] | ACTACGAATGCCCTGACGAGAAACACCAGCAG | |
| 21[88] | 18[80] | GGAATCGTCGTTTTCATCGGCATT | |
| 31[144] | 28[152] | TTGATACCGATAGTTGTACAAACT | |
| 22[39] | 25[48] | TTACCAGAGAGATGGTTTAATTTCA | 3´ - TCO/Cy5 (9nm) |
| 17[120] | 13[119] | TAATTGCTCAACCGATTGAGGGAGTACATACA | |
| 26[111] | 29[111] | GCCACCACCCTCATTGATTATACCAAGCGCGA | |
| 18[101] | 21[111] | GTAGCGCATAAATATTCATTGA | |
| 15[112] | 12[117] | TCATTAAAGGTGAATTAGATAGCCGAA | 5´ biotin |
| 6[135] | 9[143] | TCCCGACTACACCCTGAACAAAGT | |
| 26[7] | 24[16] | GCGAAAATGTTTGCCCCAGAACGAGTAGTAAAAGTTGCAG | |
| 11[8] | 11[7] | GTCATAGCATTCCACACAACATACTAATCATG | |
| 33[88] | 29[87] | AGTAACATAGGAACAACTAAAGGATCTTTGAC | |
| 33[24] | 30[8] | TTACAAACAAGTTTCCATTAAACGGGTAATGT | |
| 3[16] | 0[24] | AATATAGGGGCCTTGACGCCCTGG | |
| 31[48] | 28[56] | AGGCTTTGTTGAAAATTCAATCAT | |
| 16[79] | 19[79] | CAATTCTGCGAACGAGTCTTTTCATAATCAAA | |
| 1[16] | 0[0] | AAACGACGAGTGACTCTATGATACCGACAGTG | |
| 4[143] | 7[135] | AGCTTTCCGGCACCGCCCAATCCA | |
| 28[111] | 31[103] | AATCCGCGACCTGCTCATTGTATC | |
| 22[111] | 27[111] | CAACTAATGAACTAACCCTTGAGTAACAGTGCATCTTTTCAGGGATAG | |
| 22[7] | 20[16] | AGTGAGACTTTTTCTTTTCCGAGAGGCTTTTGTTTGCGTA | |
| 21[56] | 18[48] | AAATGTTTTTATTAGCGATTAAGA | 3´ - TCO/Cy5 (18nm/9nm) |
| 3[96] | 5[102] | TCCAAGAAGATAAGTCGCCAGTT | 3´ - Cy5 (3nm) |
| 5[120] | 2[112] | TATCGGCCGAACGCGCCTGTTTAT | |
| 24[79] | 27[71] | TGCCCCCTGCCTATTTGCGCATAG | |
| 1[24] | 3[47] | GCCAGTGCCAAGCTTTCTCAGGAGTAAGTTGGAAGGGGGA | |
| 7[144] | 4[144] | ACGATTTTTTGTTTAAATCGTAGGAATCAGCC | |
| 3[118] | 0[120] | AGTACCGCACAGGGCTTA | |
| 5[48] | 0[56] | TGGGATAGGCGGGCCTTGTGCTGCTGTAGAAACAAAATAA | |
| 31[168] | 32[152] | TACCATATCTGAATAATGGAAGGGCGCCGACA | |
| 7[16] | 4[24] | GGGCTTAAGCTACGTGATCGGCTG | |
| 30[143] | 32[120] | GGATTTTGTAAACAGCAACCATCGCCCACGCATAACCGAT | |
| 9[80] | 4[88] | AAATATTTGCATTAAACCAGAGCCTAATTTGCCGCCATTC | |
| 16[183] | 16[184] | GCTTCTGTATCCTTGAAAACATAGATAACCTT | |
| 13[120] | 8[112] | TAAAGGTGAGCTGATAAATTAATGCGGGAGAAATAAAAAC | |
| 2[39] | 7[47] | GTAACGCCGGCGGATTGACCGTAATTCATCAACGTCTGGC | |
| 1[120] | 3[143] | TGTAATTTAGGCAGAGGCATTTTCATGTTCAGAACAAGCA | |
| 4[15] | 1[15] | CACATAAATCATTTCTCTCGTCGGGTAAGCAACGGCCCTGCCATTGTA | |
| 1[80] | 3[71] | GAATTCATGTCAACCTTATGACAATGTCCCGCCCAATCAA | |
| 18[135] | 21[143] | GCGACAGAAGGCAGGTCAGACGAT | |
| 22[143] | 25[143] | CCACATCAGTTGAGATGGCGGATAAGTGCCGT | |
| 7[104] | 11[103] | AAATAAACAGGGAAGCTATTTTGTAAGGGTGA | |
| 18[39] | 23[47] | GAAAGACTGTTTTGCCAGAGGGGGACCCTCGTATTTTAAG | |
| 21[152] | 17[159] | ATATTCACTCGATAGCAGCACCGTCTCATCACCAGTAGCA | |
| 8[55] | 4[56] | TAACCAATCTTCCTGTTTACCAACCGATCGGT | |
| 22[167] | 25[183] | AGCGCAGTAGTATAGCCCGGAATAGGTGTTCG | |
| 11[128] | 15[135] | CCATCAATATGATAATTTTAAGAAAAGTAAGCATCACCGT | |
| 26[135] | 29[143] | ACCCTCAGGATTTGTATCCACAGA | |
| 5[184] | 7[175] | ATCTTCTGTTTTAGTTAATAAGGCTTATCCGGCGCGAGAA | |
| 9[120] | 6[112] | TTAACTGATGCGGGAGGTTTTGAA | 3´ - TCO/Cy5 (3nm) |
| 28[175] | 31[167] | TAACGTCAGATGAATATTAGAACC | |
| 8[47] | 11[39] | AGGAACGCCATCAAAAAAATTTTT | |
| 14[39] | 19[47] | AAATTAAGCGGAAGCAAACTCCAAGGAAGCCCCGGATTGC | |
| 4[47] | 7[39] | CTTCGCTATTACGCCAATAATTCG | 5´biotin |
| 28[87] | 24[88] | TTAGCCGGACCTTCATCAAGAGTACCGTATAA | |
| 5[112] | 3[117] | GACGACAGCCGGAAACAAACCA | 5´ - TCO/Cy5 (3nm) |
| 18[127] | 16[112] | ATCAAGTTTGCCTTTATTAGAGCTACGGTGTC | |
| 1[88] | 3[111] | GCGCACGACTTAAGTGTTTAACAACAACAATACGGGTATT | |
| 25[152] | 21[151] | TTGATATACTCTGAATTTACCGTTTGGCCTTG | |
| 8[183] | 8[184] | TGCTGATGAACTTTTTCAAATATATATGTAAA | |
| 4[79] | 7[71] | CAACTGTTGGGAAGGGGCTAACGA | |
| 2[167] | 5[183] | GTCCAGACTAGCAAGCAAATCAGATATAGTTC | |
| 20[15] | 19[7] | TTGGGCGCCAGGGTGGAGCTGCAT | |
| 30[79] | 32[56] | ATTGCGAAAAAAGGCTTTTGCGGGATCGTCACCCTCAGCA | |
| 22[103] | 25[111] | GCAGATACGATTAGGATTAGCGGG | |
| 7[112] | 4[120] | AGCCATATTATTTATCTTCTGGTG | |
| 31[112] | 28[120] | AGCTTGCTTTCGAGGTTGATAAAT | |

| | | | |
|---|---|---|---|
| 26[71] | 29[79] | AAATCAACTACACTAAAACACTCA | |
| 20[79] | 23[71] | ATGACCATAAATCAAACAGTCAGG | |
| 30[103] | 32[112] | GAATAGAATATCATTTTGCGGAACAAAGAAACATATTCGG | |
| 23[8] | 23[7] | GCTGATTGCAAGCGGTCCACGCTGGGGCAACA | |
| 12[52] | 8[56] | TTTATCATATATTTTAAATGCTCATTTTT | |
| 16[23] | 12[24] | GCGTTGCGTGAGTGAGCTAACTCATTGTTATC | |
| 30[7] | 28[16] | TCCAGTTTGGGTTGAGTGTAATACGTAATGCCAAATCAAA | |
| 10[143] | 13[143] | AACATTCAACCGTTCTGCAACATATAAAAGAA | |
| 10[39] | 13[47] | AGAGAATCAACATTATGACCCTGT | |
| 24[52] | 27[39] | ATTTTAATCATTGTGAATTACGAAAGAGG | |
| 4[175] | 7[167] | TTGAAATACCGACCGTACAAAGAA | 5´biotin |
| 27[48] | 24[53] | ACGGTGTACAGACCAGCGGAACCTATT | 5´biotin |
| 32[183] | 33[191] | TTATACTTCAAAATTATTTGCACGTGTTTGGAATCCTGAT | |
| 6[143] | 8[128] | AGCGAACCAATAAGAAAGCAGCCTTTACAGAG | |
| 10[63] | 9[55] | TTGCCTGAGAGTCTGGAAAAGCCC | |
| 1[112] | 0[96] | CGCCAACAATTGAGAATCGCCATATCCTTAGT | |
| 13[80] | 8[88] | TTATTACGGCAATAATGTAGGTAAAGATTCAATAAAATTC | |
| 14[47] | 19[39] | AATTAGCAAATAACCTATTAGATACATTTCGCAAGCAAAG | |
| 18[7] | 17[23] | GTCGTGCCCAGTCGGGAAAGAGCTTCAAAGCG | |
| 12[116] | 8[120] | CAAAGGGAGACAGTCAAATCAAGAATAAC | |
| 28[119] | 24[117] | TGTGTCGACAAGCCCAATAGGAACTTTAACGGGGT | |
| 20[151] | 16[144] | CCGCCACCCCCTCAGAGCCACCACTAGAGCCAGCAAAAAC | |
| 30[71] | 32[80] | TAATAATTAACGTTATTAATTTTAAAAGTTTGAGGCCGCT | |
| 5[103] | 6[102] | TGAGGGGACGCCTTAAATC | 3´ - TCO/Cy5 (18nm/6nm/3nm) |
| 10[71] | 15[79] | TCAGGTCAGCATGATTAAGACTCCAACATCCACGCGAGCT | |
| 12[183] | 12[184] | TCAATAGTCCTTTTTAACCTCCGGGAGAAGAG | |
| 20[47] | 23[39] | TGACTATTATAGTCAGCTTATGCG | |
| 19[16] | 16[24] | CGGCCAACGCGCGGGGCATTAATT | 5´biotin |
| 24[183] | 24[184] | ACCAAGTTAATTACCTGAGCAAAACTTTGAAT | |
| 17[80] | 12[88] | CCTTTTGATTGATTCCGAAAAGGTGGCATCAAGAGGAAAC | |
| 23[128] | 27[135] | AGGTAGAAAGATTGTATGTACTGGTAATAAGTCCATGTAC | |
| 27[144] | 24[152] | TGAGTTTCGTCACCAGTTTGATGA | |
| 18[143] | 20[128] | AATCAGTAAACCGCCAAGAACCACCACCAGAG | |
| 16[15] | 15[7] | CTCACTGCCCGCTTTCAGCCTGGG | |
| 29[184] | 30[179] | AGAAATTGTAAAACAGAAACG | |
| 18[167] | 21[183] | TGAAACCAAAACAAATAAATCCTCATTAAAAC | |
| 10[83] | 11[79] | CTACAAAGGCTAAGTAATGT | |
| 19[48] | 16[48] | ATCAAAAAGTTTGCCATAGATTTAGTTTGACC | |
| 9[152] | 5[159] | AATTGAGCGAACGCGAGGCGTTTTCCATTACCGCGCCCAA | |
| 8[79] | 11[71] | TTTTTGTTAAATCAGCAATGCCTG | |
| 4[111] | 7[103] | CAGGCAAAGCGCCATTCAGTTACA | |
| 23[144] | 20[152] | AGCGTCATACATGGCTCCTCAGAG | |
| 26[48] | 31[39] | ATTCAGTGAACAGATGACGAACTGACCAACTTTGTAGCAAC | |
| 4[183] | 4[184] | TTAATGGTTAAATAAGAATAAACAACCTAAAT | |
| 11[144] | 8[152] | GAAATAGCAATAGCTACGTCAAAA | |
| 18[73] | 23[79] | CATAGCCCCCAGACTGGATAGCGTCCTTACGAGGACGTTGGG | |
| 14[135] | 17[143] | GCGACATTGAATATAATGCTGTAG | |
| 25[184] | 27[175] | CCTGATTGCAATAACGGATATCACCGTACTCACAGTACCT | |
| 13[152] | 8[144] | ACACCACGAATAATAAGAGCAAGACAGAGGGTATGAAAAT | |
| 10[47] | 15[39] | AGCAAACAAGAACCCTTTCAACGCAAGGATAAAAATGGTC | |
| 19[104] | 23[103] | GGAACCGCCTCAAATGCTTTAAACAATAAAAC | |
| 26[39] | 29[47] | ATAAGGCTGGCACCAACCTAAAAC | |
| 1[48] | 3[39] | AAGCCAGGCCCCGCTTCTAATCTATTTACGCTGCTGGCGA | |
| 0[151] | 3[135] | AGTATAAAGCCAACGCTCAACAGTTCATCGAG | |
| 27[16] | 24[24] | TCCGAAATCGGCAAAACCTGAGAG | |
| 30[178] | 33[183] | TCTTTCCAGACGTTAGTAAAGATGATGGCAATTCATCAATATA | |
| 1[152] | 1[183] | AATAAGAGAATATAAAGTACCGACAAAAGATT | |
| 5[80] | 0[88] | TCGTAACCAGGCTGCGTGTCTTTCCTTATCATGCTGAATT | |
| 0[191] | 3[175] | AAGCCTGTACTAGAAACCGGAATCATAGTAAAGTAATTCTTAAGGCGT | |
| 6[101] | 13[111] | AAGATTAACGTTTAGCTATAGCAAACGTAGAAAA | |
| 14[143] | 19[135] | ACAAAAGGCACCGACTATGTTTTAAATATGCAACCCTCAG | |
| 0[183] | 3[167] | TTAGTATCATATGCGTTATACAAAGTGATAAA | |
| 16[175] | 19[167] | AAATCGTCGCTATTAAATTTGAAT | |
| 2[71] | 5[79] | TCCTAATTGGTGTAGATGGGCGCA | |
| 30[135] | 32[144] | CTAAACAAGGAGCGGAATTATCATCATATTCCATGACAAC | |
| 10[167] | 13[183] | TTAAGCCCGAATAAGTTTATTTTGTCACAATT | |
| 6[39] | 9[47] | CATTAAATCCCGGTTGATAATCAG | |
| 7[8] | 7[7] | AGTAAACATAAAGACGGAGGATCCGTGTAATG | |
| 11[104] | 12[96] | GAAAGGCCTTACCAGAAGGAAACC | |
| 29[120] | 25[119] | ACAACGGAAACCGCCACCCTCAGAGTTTTGCT | |
| 12[175] | 15[167] | GAATTTATCAAAATCATTAATTTT | |
| 30[111] | 32[88] | CGGAGTGAGGTTTATCTCGCTGAGGCTTGCAGGGAGTTAA | |

| | | | |
|---|---|---|---|
| 33[152] | 28[144] | TGATTATCATGAATTTTCTGTATGCAGCCCTCACAACGCC | |
| 17[102] | 18[102] | TGCGGATGGCGCGTCAGACT | 3´ - TCO/Cy5 (18nm) |
| 24[116] | 20[112] | CAGTGGGAACAACATTATTACCCGCCGCCAGCATCCC | |
| 12[79] | 15[71] | AACGGAATACCCAAAAATTTGGGG | |
| 31[8] | 31[7] | AGTCCACTGAAAAACCGTCTATCAGGAACAAG | |
| 28[79] | 31[71] | AACGAGGCGCAGACGGCTCCAAAA | |
| 6[15] | 4[16] | CCGTCCCTCCTGGTTGCCGAACTCTGAGGATTCTCCGTGGACGCATTT | |
| 15[144] | 12[152] | TGAGCCATTTGGGAATTCTTACCG | |
| 9[88] | 6[84] | AAATTGTAAGTTGCTATTTT | 3´ - TCO/Cy5 (6nm) |
| 17[48] | 13[63] | CAGGTCAGGATTAGAGTACAGGCAAGGCAAAGAATACTTTTGCGGCTG | |
| 20[183] | 20[184] | AAATTAATTTAATGGAAACAGTACAGAAAACA | |
| 15[96] | 19[103] | TTCTGACGGAAATTATTGGAAGTTTCATTCCACCACCACC | |
| 33[120] | 29[119] | CACCAGAACTTTCAACAGTTTCAGAACAAAGT | |
| 14[167] | 17[183] | TCATATGGCCATTACCATTAGCAAGGCCGTAT | |
| 6[71] | 9[79] | AATTTTATGGAAGATTGTATAAGC | |
| 10[7] | 8[16] | CGAATTCGCCGGGTACCGATAGCATGTCAATCTACCTCGA | |
| 1[56] | 3[79] | GTGGATGTTCTTCTAAGTGGTTGTATATCCCATAATCGGC | |

**Single molecule DNA origami surface preparation.** For the preparation of DNA - Origami single-molecule surfaces, 8 chambered cover glass systems with high performance cover glass (Cellvis, C8-1.5H-N) were used. The surfaces were washed once with PBS (Sigma-Aldrich, D8537-500ML) prior treatment with 2% Hellmanex (Hellma, 9-307-011-4-507) for 1 hour. After washing the chambers three times with PBS (Sigma-Aldrich, D8537-500ML), the surfaces were incubated with 1 M KOH (Fulka, 06005) for 20 min. After alkaline treatment, the chambers were washed with PBS (Sigma-Aldrich, D8537-500ML). Afterwards, the surfaces were incubated with 10 % polyethylenglycol 400 (Fulka, 81170) over night at 4 °C. Afterwards, the surfaces were rinsed 3 times with PBS (Sigma-Aldrich, D8537-500ML) before incubating the chambers with 0.5 g/l BSA-Biotin (ThermoFisher, 29130) in PBS) overnight at 4 °C. In the following, the chambers were washed three times with PBS (Sigma-Aldrich, D8537-500ML) before incubation with 0.5 g/l Neutravidin (ThermoFisher, 31050) in PBS (Sigma-Aldrich, D8537-500ML) for 20 min. The surfaces were washed three times with PBS (Sigma-Aldrich, D8537-500ML) and incubated with purified DNA – Origami solution, 1:5 diluted in PBS (Sigma-Aldrich, D8537-500ML) + 50 mM MgCl2 (AppliChem, A4425,0500) for 10 min. The prepared samples were washed at least three times in PBS (Sigma-Aldrich, D8537-500ML) + 50 mM MgCl2 (AppliChem, A4425,0500) prior to imaging.

**Cell culture.** HEK-293-T cells (German Collection of Microorganisms and Cell Cultures, Braunschweig, Germany; #ACC635) were maintained in T25-culture flasks (Thermo Fisher, Cat. Nr. 156340) in Dulbeccos´s Modified Eagle´s Medium (DMEM, Sigma-Aldrich, #D5796) supplemented with 10 % FCS (Sigma-Aldrich, #F7524), and 1 % Pen-Strep (Sigma-Aldrich, #P4333)

**Plasmid constructs.** All plasmids were amplified by transformation to E. coli XL1 – Blue followed MIDI-prep DNA isolation and sequencing (Nucleobond®, Xtra Midi, Macherey & Nagel, #740410). The plasmid for the expression of clickable α2 subunit of the GABA-A receptor was obtained from Addgene (Addgene # 49169) (*59*). The superecliptic pHluorin tag was removed by introducing a XhoI restriction site upstream of the GABA-A coding sequence and subsequent cutting with XhoI-XhoI. The plasmids for the expression of the GABA-AR β1 and γ2 subunits were kindly provided by Andrea

Barberis and described previously (*60,61*). The plasmid for the expression of clickable GluK2 was a kind gift from Peter Seeburg (*62*). The amber stop mutants of GluK2, GABA-AR α2 and GABA-AR γ2 subunits were generated by introducing a TAG stop codon via PCR-based site-directed mutagenesis of the vectors using custom designed primers (Sigma) and Q5 High-Fidelity DNA Polymerase (New England BioLabs). The plasmid for the expression of the tRNA/aminoacyl transferase pair (pCMV tRNAPyl/NESPylRSAF, herein termed PylRS/tRNAPyl) was kindly provided by Edward Lemke (*63*). The plasmid for the expression of the tRNA/aminoacyl transferase pair (pNEU-hMbPylRS-4xU6M15, herein termed PylRS/4xtRNAPyl) was a gift from Irene Coin (Addgene, #105830) (*64*).

**Transfection of HEK293T cells.** Transfection of HEK293T cells was carried out using the JetPrime Transfection Reagent (Polypus, #114-01) according to manufacturer instructions. HEK293T cells were seeded on 4-well Lab-Tek II chambered glass slides (Nunc, cat. no. 155409) coated with 0.5 mg/ml poly-D-Lysine (Sigma-Aldrich, #P6407) the day before transfection. At 70-85% confluency the cells were transfected. Transfection of GluK2 receptors was carried out with 500 ng GluK2 and 500 ng pCMV NES-PylRSAF/tRNAPyl per well. GABA-A receptor subunits were transfected at the following ratio with a total amount of 1750 ng DNA per well: 500 ng α2 subunit, 500 ng β1 subunit, 250 ng γ2 subunit and 500 ng pCMV NES-PylRSAF/tRNAPyl. Additionally, the cells were fed the unnatural amino acid TCO*-A (SiChem, SC-8008) supplemented to the cell media. Therefore, the TCO*-A was diluted 1:4 with 1M HEPES (pH 8.0) and added at a final concentration of 250 μM to the cells. Transfected cells were maintained in an incubator with 5% CO2 at 37°C for 24h (GluK2) or 48h (GABA-AR) depending on transfected constructs and subsequently labeled with fluorophores.

**Bioorthogonal click labeling of receptors.** Transfected HEK-293T expressing the TCO*-A modified GluK2, or GABA-A α2 or GABA-A γ2 receptor subunits were labeled with 3 μM tetrazine coupled fluorophores H-Tet-Cy5 (Jena Bioscience, #CLK-015-05) in cell growth medium for 60 min on ice. Then, cells were washed three times with ice-cold PBS. Next, fixation was carried out with 4 % formaldehyde and 0.25 % glutaraldehyde for 15 minutes at room temperature. Following fixation, cells were again washed three times with PBS and subsequently imaged at the *d*STORM setup.

***d*STORM and DNA-PAINT imaging.** Super-resolution imaging was performed using an inverted wide-field fluorescence microscope (IX-71; Olympus). For excitation of Cy5, a 641 nm diode laser (Cube 640-100C, Coherent), in combination with a clean-up filter (Laser Clean-up filter 640/10, Chroma) was used. The laser beam was focused onto the back focal plane of the oil-immersion objective (60×, NA 1.45; Olympus). Emission light was separated from the illumination light using a dichroic mirror (HC 560/659; Semrock) and spectrally filtered by a bandpass filter (FF01-679/41-25, Semrock). Images were recorded with an electron-multiplying CCD camera chip (iXon DU-897; Andor). Pixel size for data analysis was measured to 128 nm. For *d*STORM measurement, 120,000 images with an exposure time of 5 ms (frame rate 200 Hz) and irradiation intensity of ~5 kW $cm^{-2}$ were recorded. Single-molecule surfaces were imaged by EPI illumination, whereas prepared cells were imaged by TIRF illumination. *d*STORM experiments were performed in PBS-based photoswitching buffer containing

100 mM β-mercaptoethylamine (MEA, Sigma-Aldrich) and 50 mM $MgCl_2$ (AppliChem, A4425,0500) for DNA – Origami measurements, or without MgCl2 for receptor imaging, adjusted to pH 7.6. For each DNA - PAINT measurement, 18,000 images with an exposure time of 100 ms (frame rate 10 Hz) were recorded. Single-molecule DNA - Origami surfaces were imaged by total internal reflection illumination, excited with a 561 nm diode laser (Genesis MX561-500 STM, Coherent) at an irradiation intensity of ~1.5 kW $cm^{-2}$ in combination with a clean-up filter (Laser Clean-up filter 561/14, Chroma). Emission light was separated from the illumination light using a dichroic mirror (FF403/497/574-Di01; Semrock) and spectrally filtered by a bandpass filter (BrightLineHC-607/70, Semrock). DNA - PAINT experiments were performed at 5 nM imager – strand concentration (5´-3´: CTA GAT GTA T, biomers.net), 5´ - modified with Cy3B, in PBS-based buffer containing 5 mM TRIS (Merck, 1.08382.2500), 50 mM MgCl2 (AppliChem, A4425,0500), 1 mM EDTA (Sigma, E1644-250G) and 0.05% Tween20 (ThermoFisher, 28320) adjusted to pH 7.6. All SMLM results were analyzed with rapidSTORM3.3 (*65*) and the highly resolved pictures were reconstructed with ThunderSTORM (*66*). The localization precisions were calculated according to Mortensen *et al.* (*67*). For photoswitching fingerprint analysis only fluorescent spots containing more than 500 (*d*STORM) / 6000 (DNA-PAINT) photons per frame were analyzed. To estimate the number of localizations per fluorophore, the tracking function (Kalman filter) of rapidSTORM3.3 was used (*63*). Fluorescent spots were tracked over the whole image stack (120,000 frames for *d*STORM and 18,000 frames for DNA-PAINT) within a tracking radius of 200 nm. The information was saved as tracked localization file. A custom written python script was used to calculate the number of frames of consecutive localizations per spot (on-time) as well as the number of frames between on-time events of the same fluorescent spot within the defined tracking radius (off-time). In addition, also the average number of photons detected per frame as well as the number of on-time events per tracked spot was calculated.

**Fluorescence lifetime intensity trajectories.** All fluorescence lifetime measurements concerning single-molecule trajectories and photon antibunching measurements were performed on a MicroTime200 (PicoQuant, Berlin, Germany) time-resolved confocal fluorescence microscope setup consisting of a FLIMbee galvo scanner (PicoQuant, Berlin, Germany), an Olympus IX83 microscope including an oil-immersion objective (60×, NA 1.45; Olympus), 2 single photon avalanche photodiodes (SPAD) (Excelitas Technologies, 75154 K3, 75154 L6) and a TimeHarp300 dual channel board. For pulsed excitation a white-light laser (NKT photonics, superK extreme) was coupled into the MicroTime200 system via a glass fiber (NKT photonics, SuperK FD PM, A502-010-110). A 100 µm pinhole was used for all measurements. The emission light was split onto the SPADs using a 50:50 beamsplitter (PicoQuant, Berlin, Germany). To filter out after glow effects of the SPADs used as well as scattered and reflected light, 2 identical bandpass filters (ET700/75 M, Semrock, 294808) were installed in front of the SPADs. The measurements were performed and analyzed with the SymPhoTime64 software (PicoQuant, Berlin, Germany). Measurements were performed with an irradiation intensity of ~ 0.5 – 2.5 kW $cm^{-2}$ in T3 mode with 25 ps time-resolution, whereas all photon antibunching measurements were performed in T2 mode. For photon antibunching experiments, the Sync cable was disconnected and replaced by the SPAD 2 cable. For analyzing the fluorescence lifetime of the trajectories, the decay parameters were determined by least-squares deconvolution, and their quality was judged by the reduced $\chi^2$ values and the randomness of the weighted residuals ($\chi^2$ = ~1). In the case that a monoexponential model was not adequate to describe the measured decay, a multiex-ponential model was used to fit the decay ($\tau_{av} = \tau_1 a_1 + \tau_1 a_1$). For reference structures and 18 nm DNA-origamis we measured a monoexponential fluorescence decay.

**Photon antibunching measurements.** Photon antibunching experiments take advantage of the fact that the probability of emitting two consecutive photons drops to zero for a single emitter for time intervals shorter than the excited-state lifetime. After photon emission, a molecule must be re-excited and wait, on average, one fluorescence lifetime before another photon can be emitted. For sufficiently short laser pulses the number of photon-pairs detected per laser pulse in photon antibunching experiments can be used to determine whether the emission is from one or more independently emitting quantum systems. As expected for *d*STORM experiments where only a single fluorophore is expected to reside in the on-state per DNA origami, the ratio of the number of photon pairs detected in the central peak at delay time zero to the average number in the lateral peaks in the interphoton-time (coincidence) histograms is <0.20 demonstrating the presence of a single emitter in the confocal laser focus with low background contributions. This result shows that although increased photoactivation at interfluorophore distances of <10 nm transfers fluorophores from the off- to the on-state the probability for two fluorophores residing simultaneously in the on-state showing independent fluorescence emission is negligible. Even if two fluorophores are simultaneously in the on-state, other energy transfer processes such as homo energy transfer and single-singlet annihilation can occur so that the on-state is dominated by the emission of a single fluorophore (*25-28*). The data in the interphoton time histograms can be quantified for the purpose of determining the number of independent emitters by determining the ratio of the number of photons in the central peak, $N_c$, to the average number in the neighboring lateral peaks, $N_{l,av}$. Ensemble antibunching measurements show that the number of photon pairs detected in the neighboring peaks decreases at large interphoton times but is nearly constant for very short times, i.e., in the first neighboring peaks. For determination of $N_{l,av}$, we used the average number of events in the nearest 8 peaks, 4 to each side of the zero-time peak.

**Time-correlated single photon counting (TCSPC).** Measurements take place in a 0.3 mm path-length fluorescence cuvette (Hellma, 105.251-QS) on a FluoTime 200 time-resolved spectrometer (PicoQuant, Berlin, Germany) in combination with a pulsed diode laser (635 nm) as the excitation source with a SepiaII module (PicoQuant, Berlin, Germany), a PicoHarp300 TCSPC module and picosecond event timer (PicoQuant, Berlin, Germany) (80 MHZ, 50 ps pulse length, 8 ps resolution, 10.000 photons in the maximum channel). The results were analyzed with the FluoFit 4.4.0.1 software (PicoQuant, Berlin, Germany). To exclude polarization effects, fluorescence was observed under the magic angle (54.7°). The decay parameters were determined by least-square deconvolution, and their quality was judged by the reduced χ 2 values.

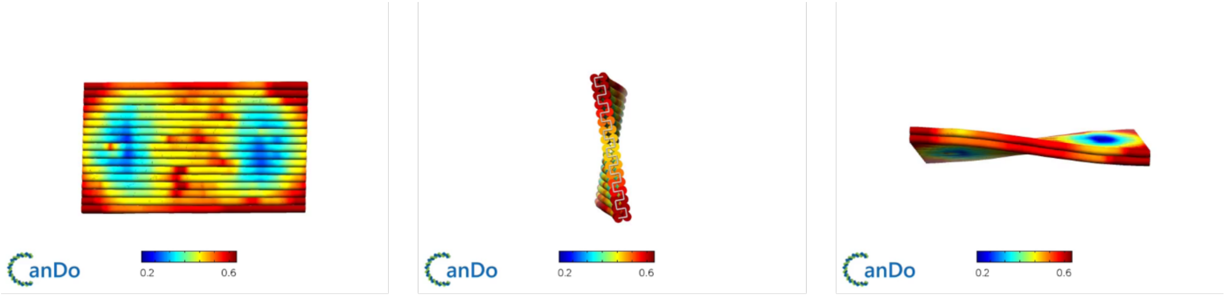


**Figure S1. DNA origami stability calculation.** The stability of the designed rectangle DNA origami structures were calculated using the open source software CanDO (*57,58*) with standard DNA parameters. Heatmap in nm.

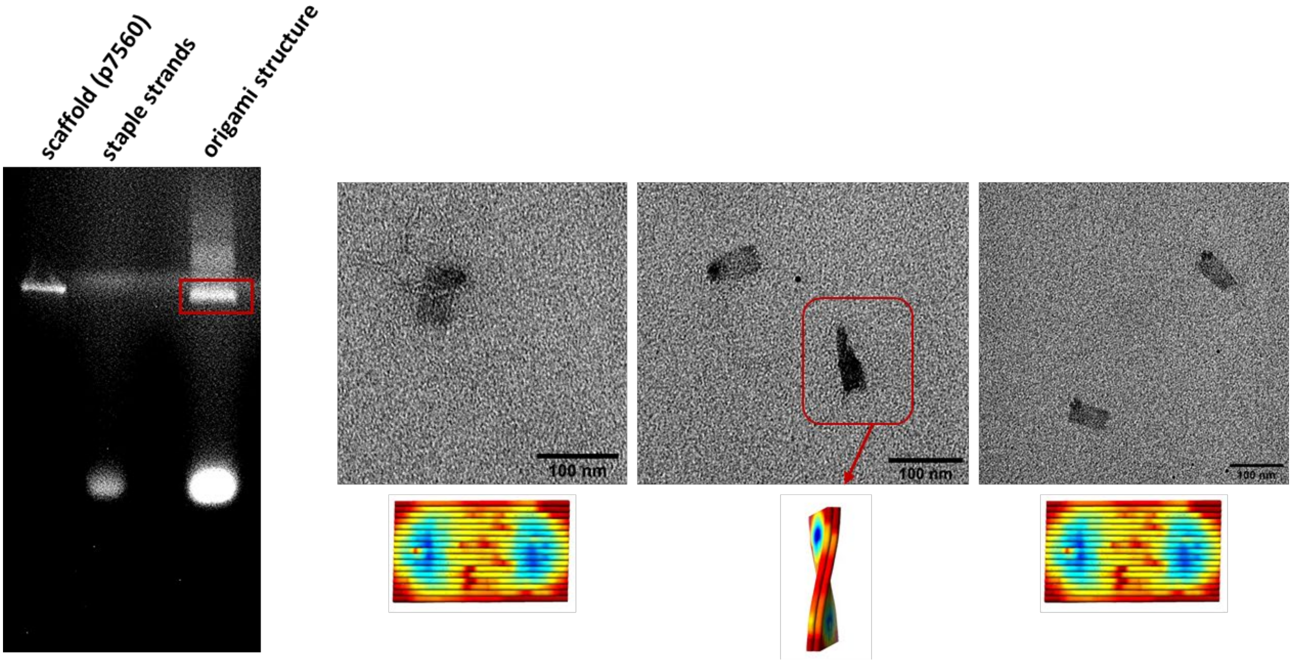


**Figure S2**. **DNA origami quality control.** Hybridized DNA origami structures were purified using 1.5% agarose gel. The marked band was cut out and the shape of the DNA origami structures were analyzed using electron microscopy.

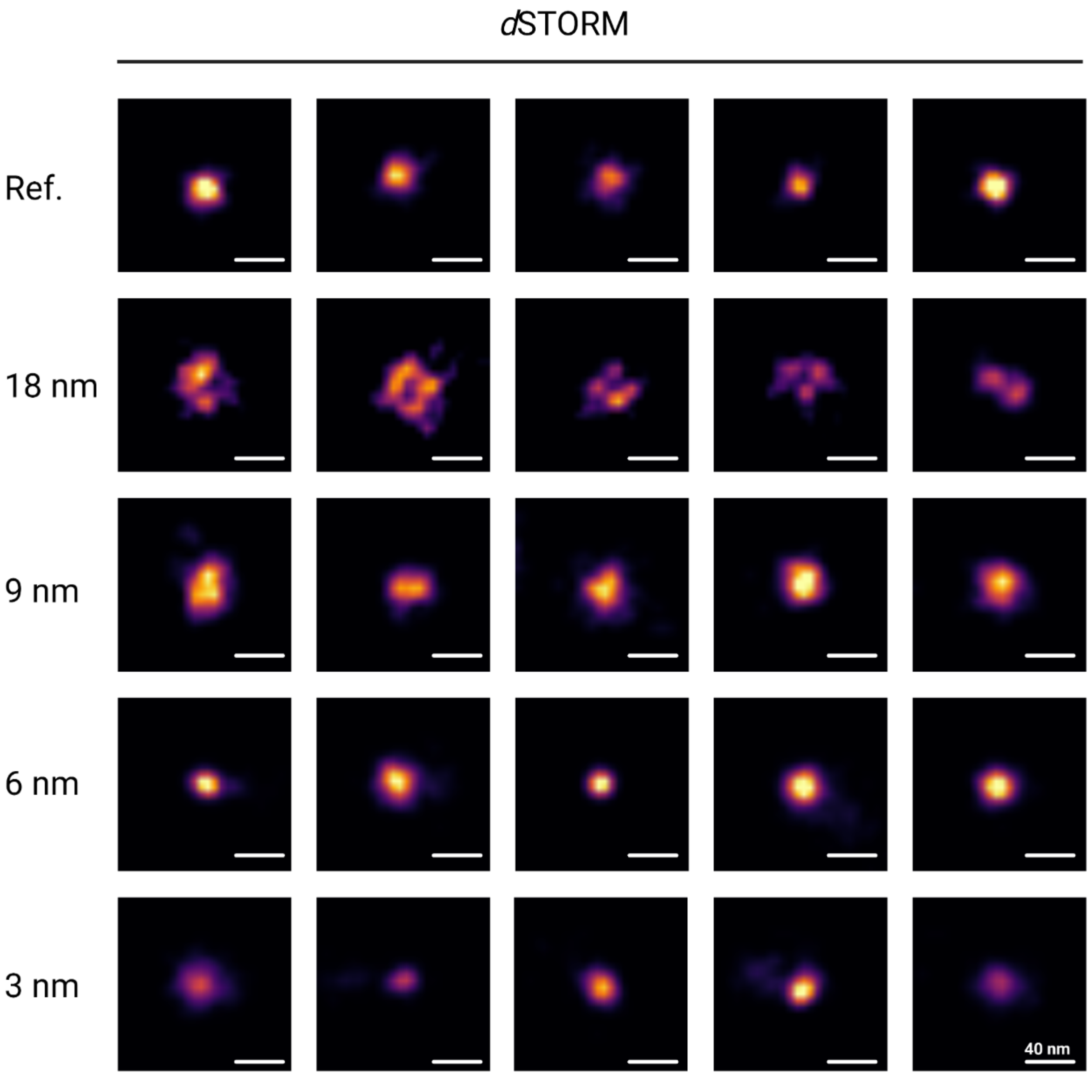


**Figure S3**. Example *d*STORM images of DNA origami with one (Ref.) or four Cy5 dyes with interfluorophore distances of 18, 9, 6, and 3 nm.

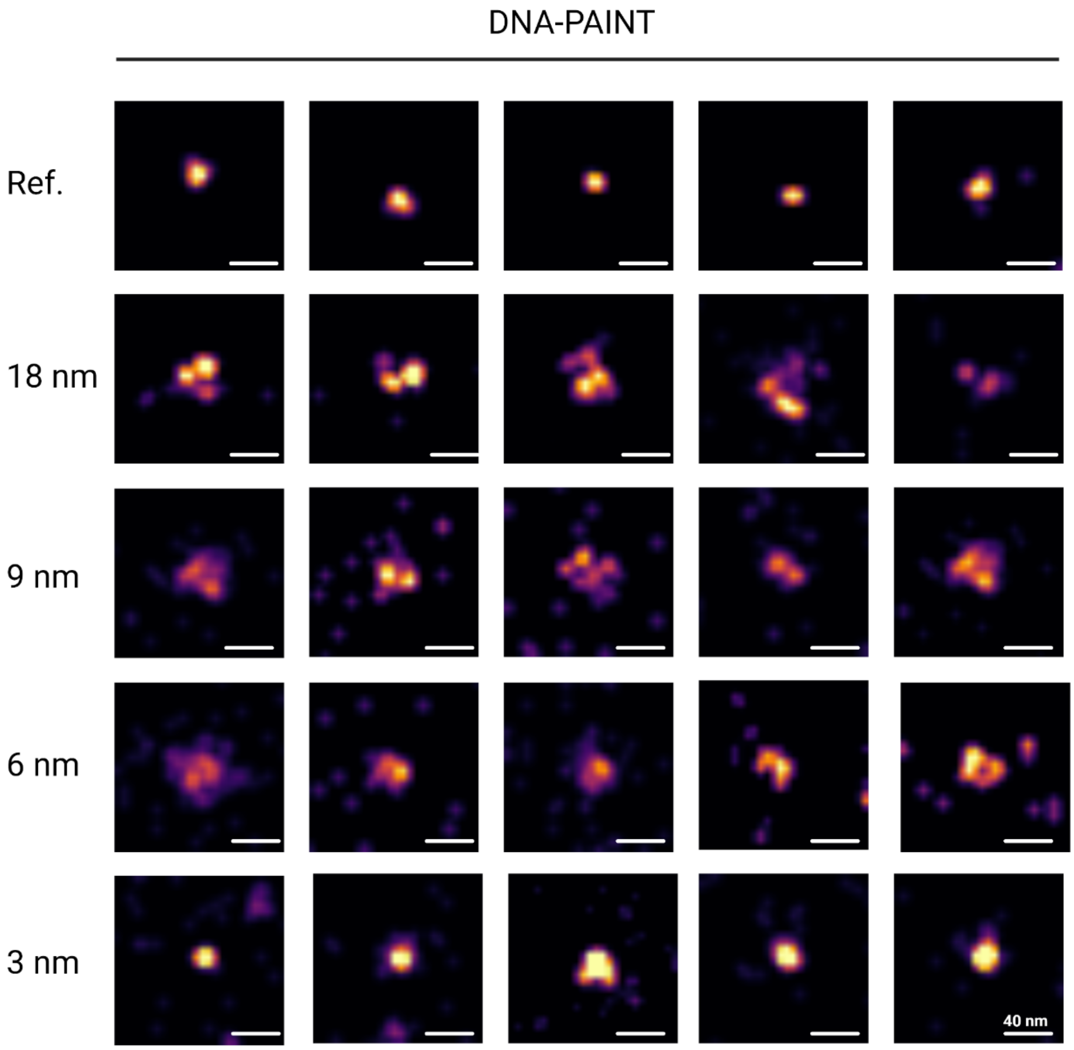


**Figure S4**. Example DNA-PAINT images of DNA origami with one (Ref.) or four docking strands separated by 18, 9, 6, and 3 nm.

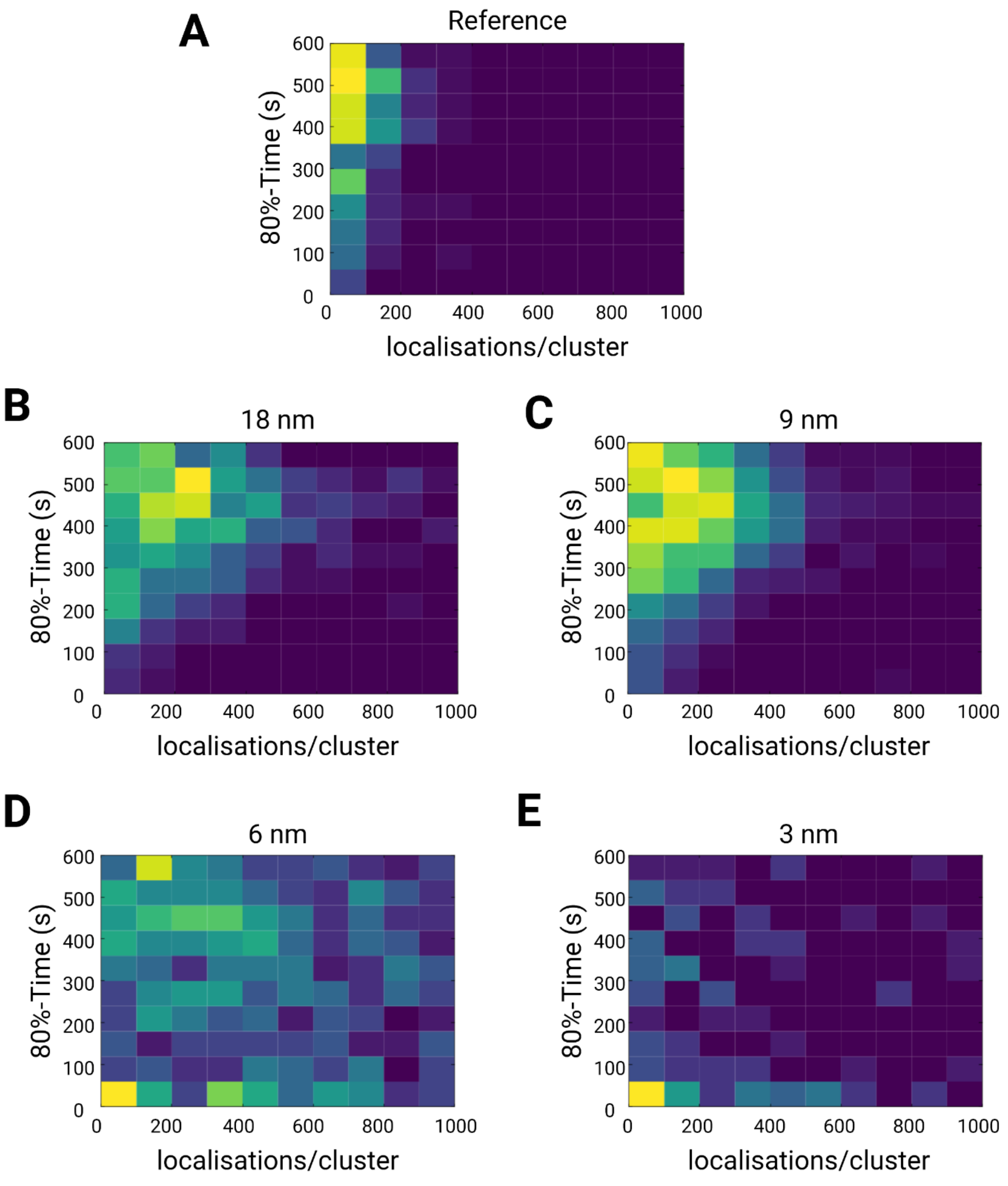


**Figure S5**. Bivariate histograms of detected localizations and the times after which 80% of all localizations were detected per individual DNA origami. The histograms clearly show that for shorter interfluorophore distances of 3 and 6 nm 80% of all localizations are detected during the first minute.

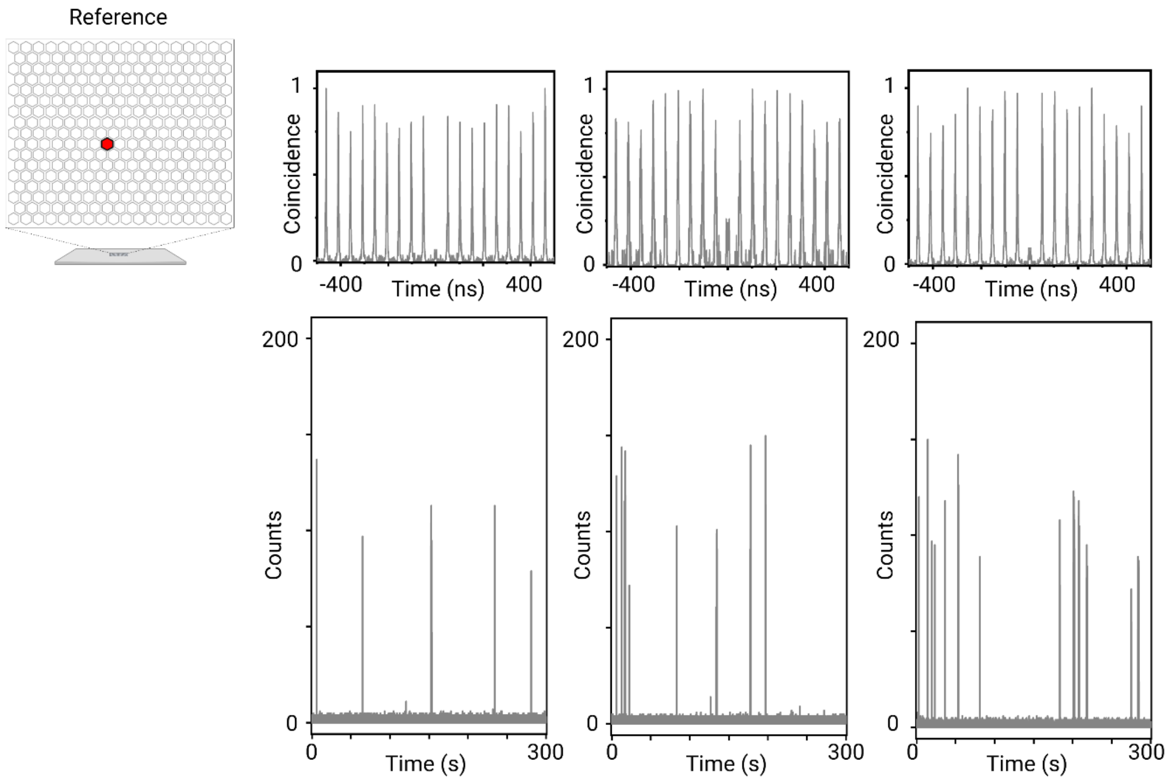


**Figure S6**. Fluorescence trajectories and corresponding normalized interphoton time (coincidence) histogram measured for the entire trajectory of singly Cy5-labeled reference DNA origami measured by single-molecule sensitive confocal fluorescence microscopy in photoswitching buffer excited at 640 nm with 2.5 kW $cm^{-2}$ (1 ms binning).

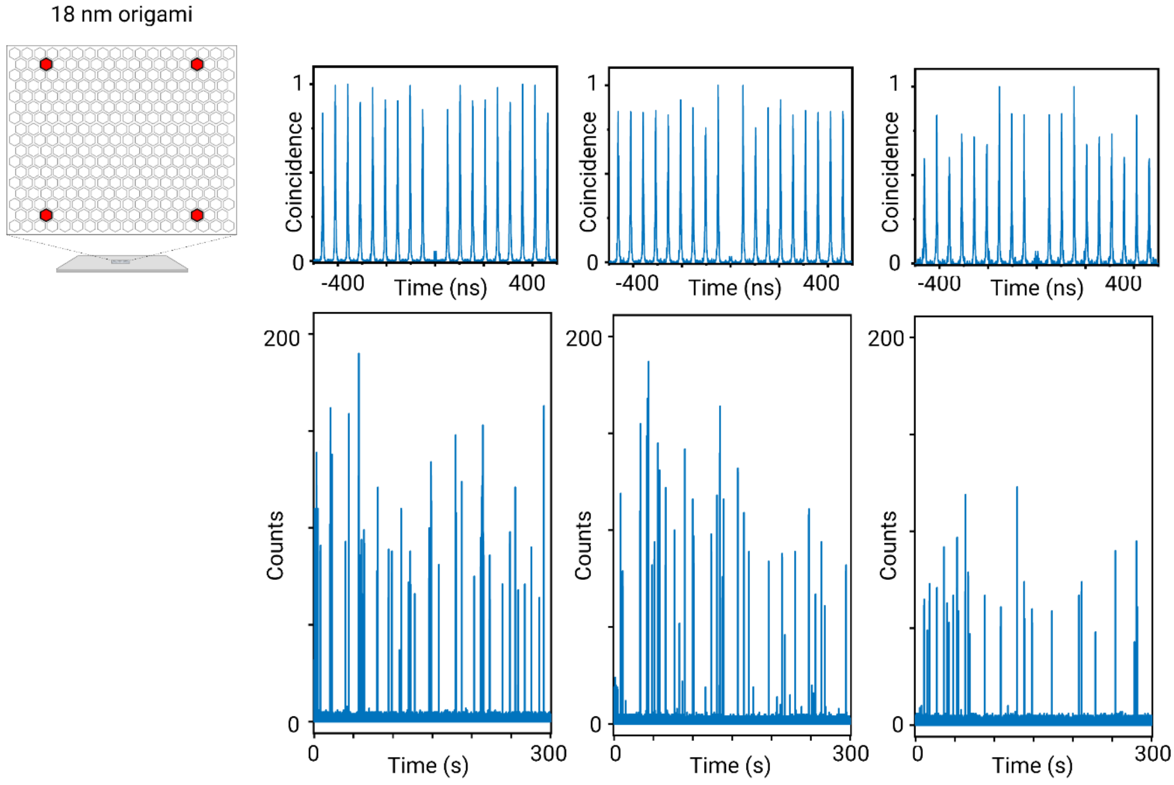


**Figure S7**. Fluorescence trajectories and corresponding normalized interphoton time (coincidence) histogram measured for the entire trajectory of 18 nm DNA origami measured by single-molecule sensitive confocal fluorescence microscopy in photoswitching buffer excited at 640 nm with 2.5 kW $cm^{-2}$ (1 ms binning).

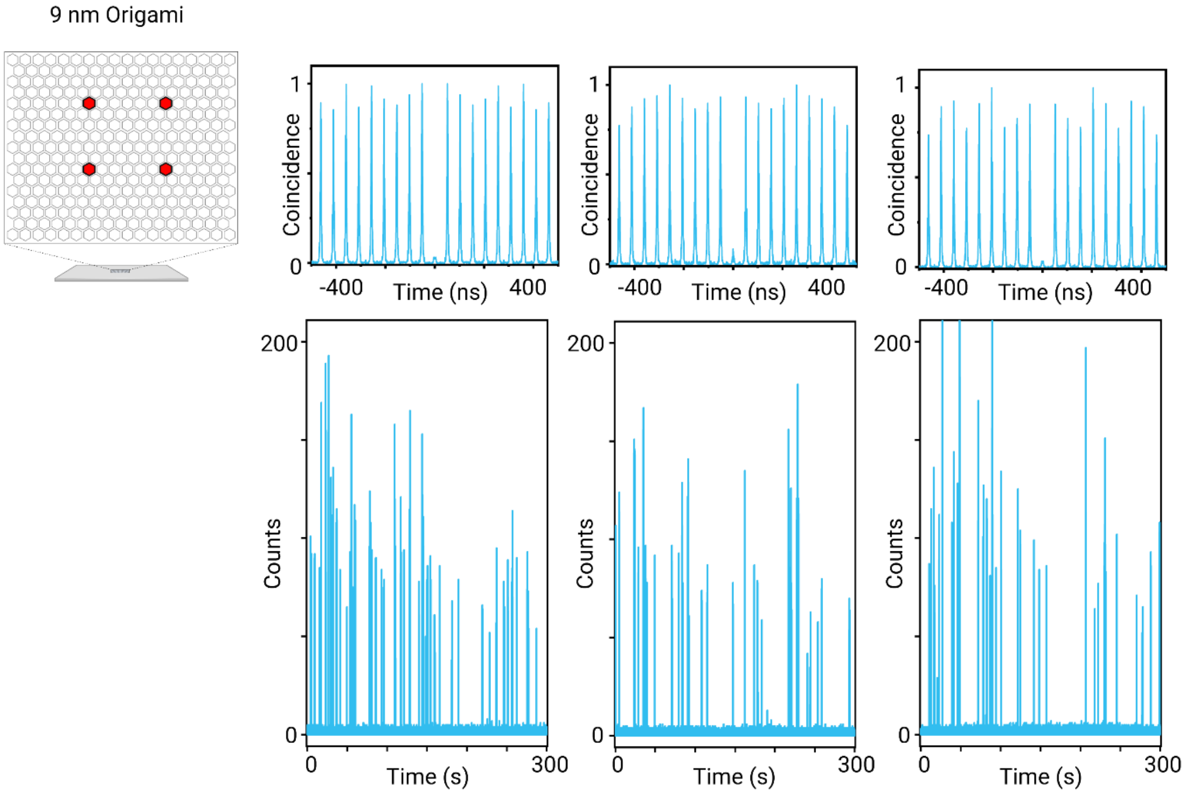


**Figure S8**. Fluorescence trajectories and corresponding normalized interphoton time (coincidence) histogram measured for the entire trajectory of 9 nm DNA origami measured by single-molecule sensitive confocal fluorescence microscopy in photoswitching buffer excited at 640 nm with 2.5 kW cm$^{-2}$ (1 ms binning).

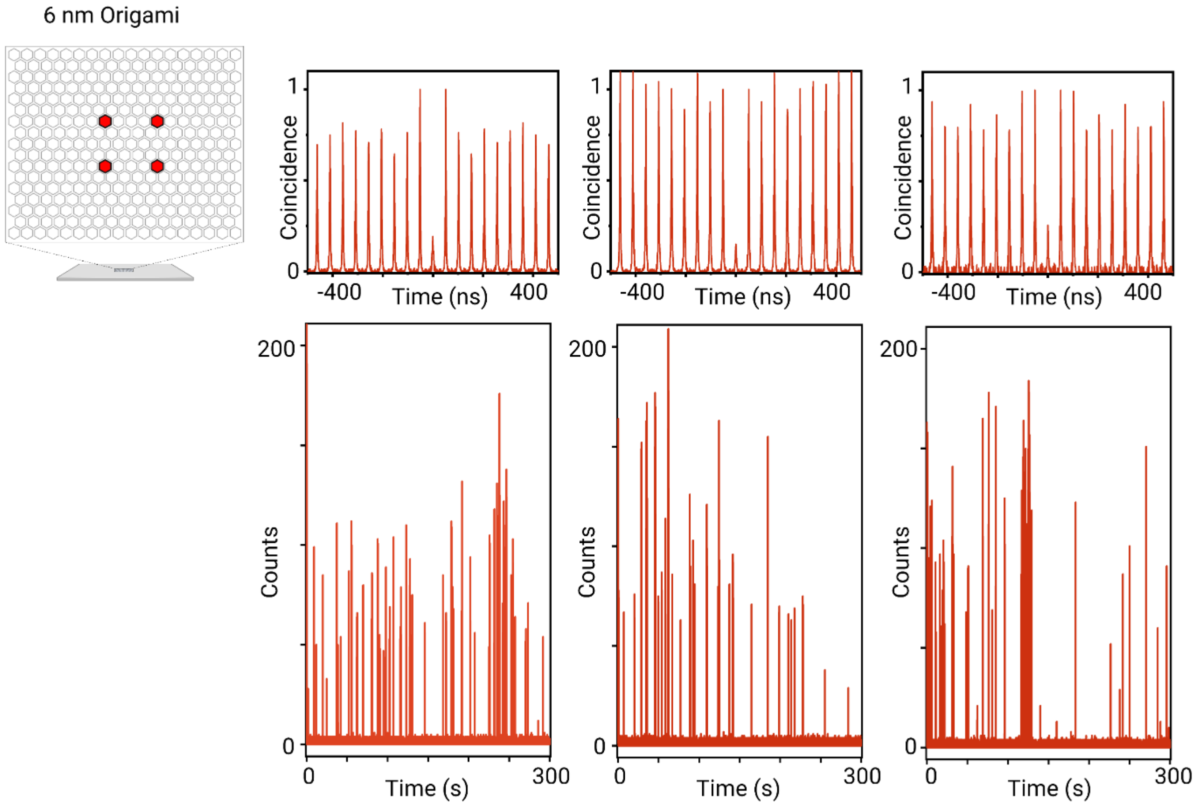


**Figure S9**. Fluorescence trajectories and corresponding normalized interphoton time (coincidence) histogram measured for the entire trajectory of 6 nm DNA origami measured by single-molecule sensitive confocal fluorescence microscopy in photoswitching buffer excited at 640 nm with 2.5 kW $cm^{-2}$ (1 ms binning).

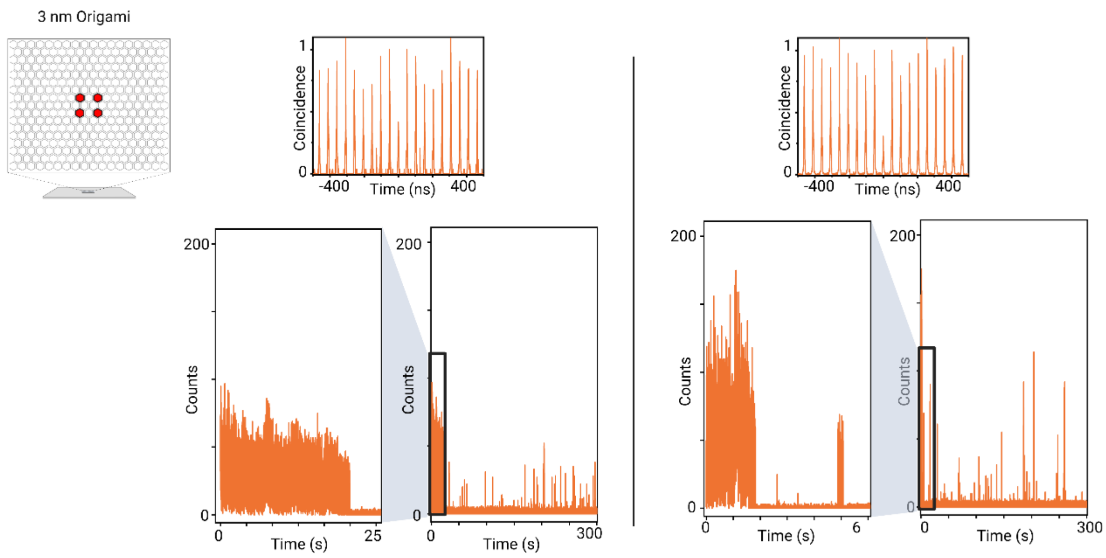


**Figure S10**. Fluorescence trajectories and corresponding normalized interphoton time (coincidence) histogram measured for the entire trajectory of 3 nm DNA origami measured by single-molecule sensitive confocal fluorescence microscopy in photoswitching buffer excited at 640 nm with 2.5 kW $cm^{-2}$ (1 ms binning).

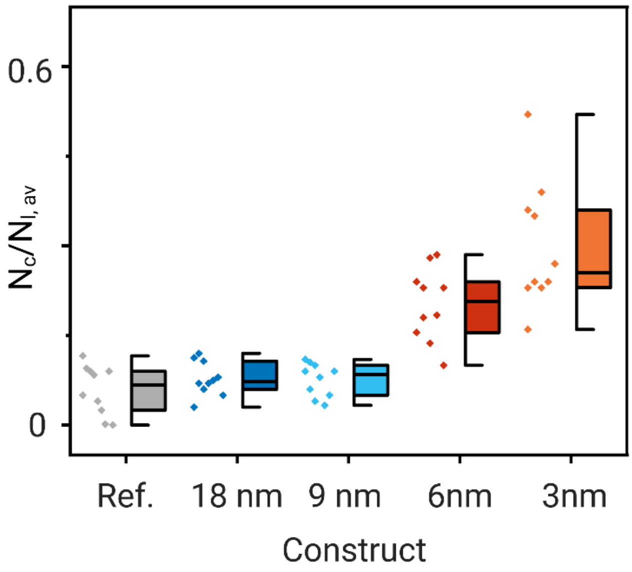


**Figure S11**. $N_c/N_{I,av}$ ratios measured for n=10 single-molecule trajectories of the different DNA origami constructs in photoswitching buffer.

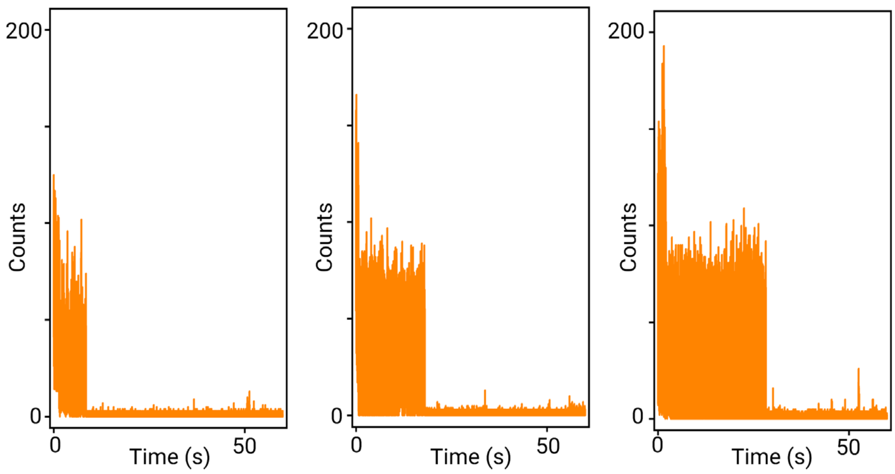


**Figure S12**. Fluorescence trajectories of 3 nm DNA origami in PBS, pH 7.6 containing 1 mM trolox/troloxquinone and an oxygen scavenging system measured by single-molecule sensitive confocal fluorescence microscopy excited at 640 nm with 2.5 kW $cm^{-2}$ (1 ms binning).

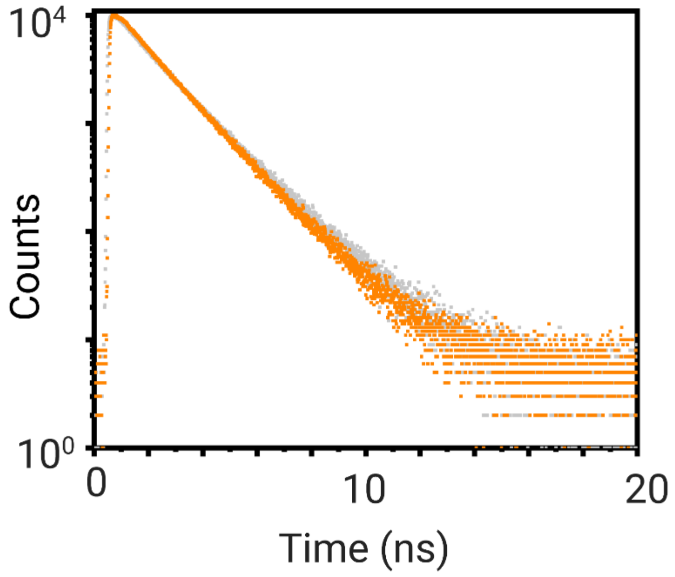


**Figure S13**. Ensemble fluorescence decays of oligonucleotides labeled with one (5´-3´: TACGATTCGATTACGTTACCATTAGCATTGCATTAGCTTATAT-*Cy5*) and four (5´-3´: TACGATTCGATT-*Cy5*ACGTTACCATT-*Cy5*AGCATTGCATT-*Cy5*AGCTTATAT-*Cy5*) dyes measured by TCSP in PBS, pH 7.6.

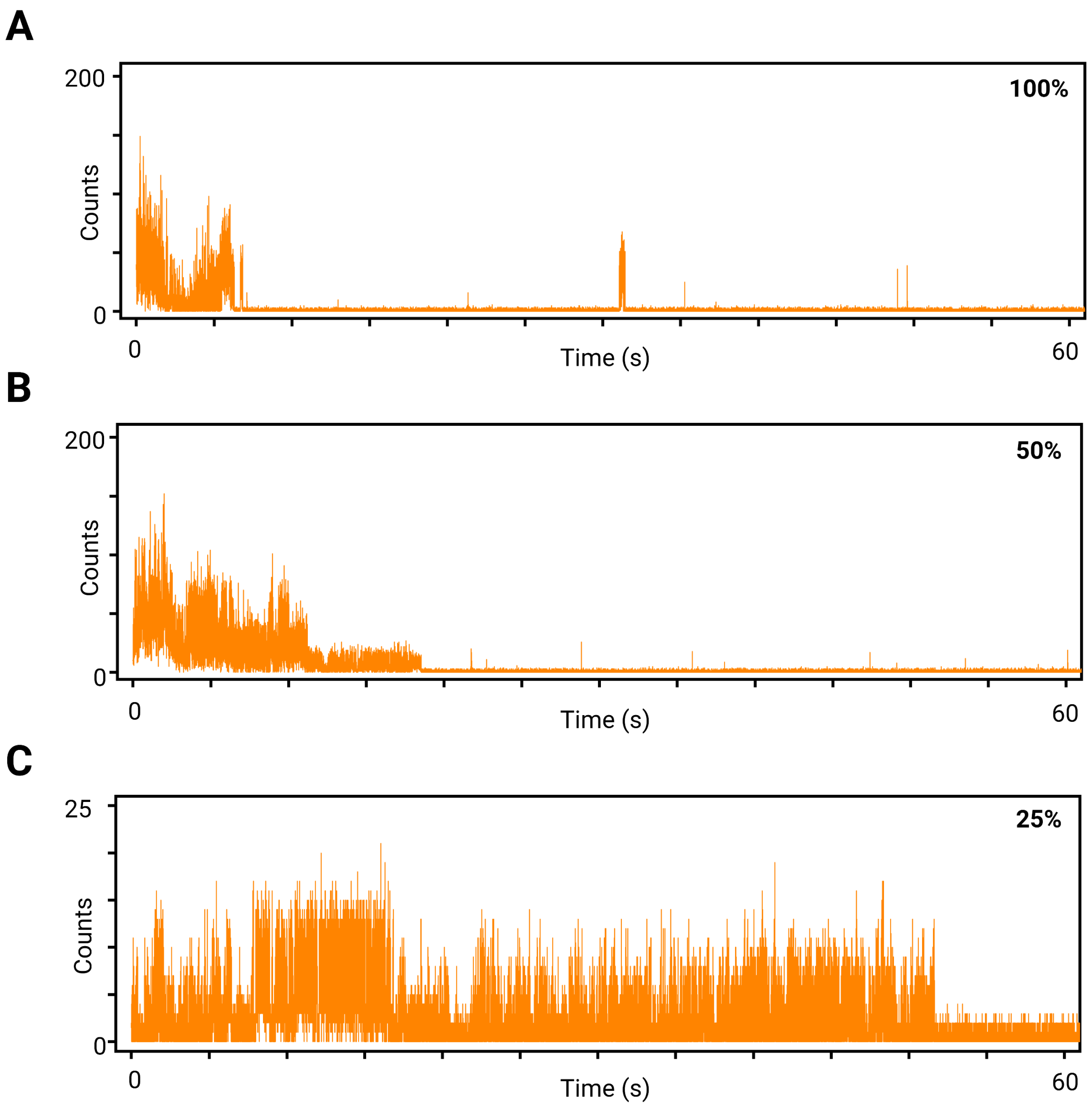


**Figure S14.** Fluorescence trajectories of 3 nm DNA measured by single-molecule sensitive confocal fluorescence microscopy in photoswitching buffer excited at 640 nm with different irradiation intensities. (**A**) 100% laser power. (**B**) 50% laser power. (**C**) 25% laser power. (1 ms binning). 100%: 2.5 kW $cm^{-2}$.

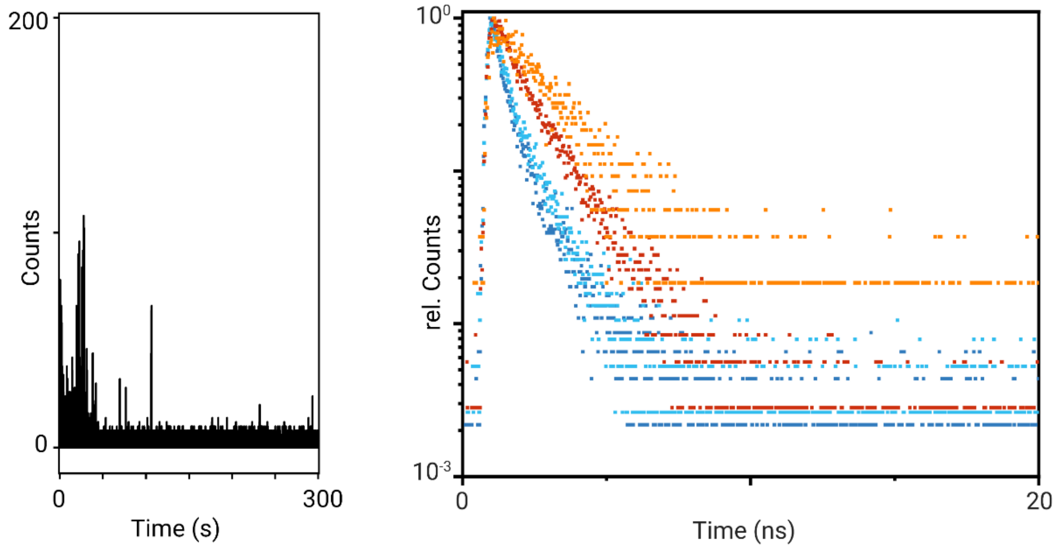


**Figure S15**. Fluorescence trajectory of a 3 nm DNA origami in photoswitching buffer and corresponding fluorescence decays recorded at different times indicate that the lifetime increases with time due to stepwise photobleaching of fluorophores and corresponding lower energy transfer efficiency (dark blue decay: 0-2 s; light blue decay: 2-25 s; red decay: 25-28 s; orange decay: 25-100 s).

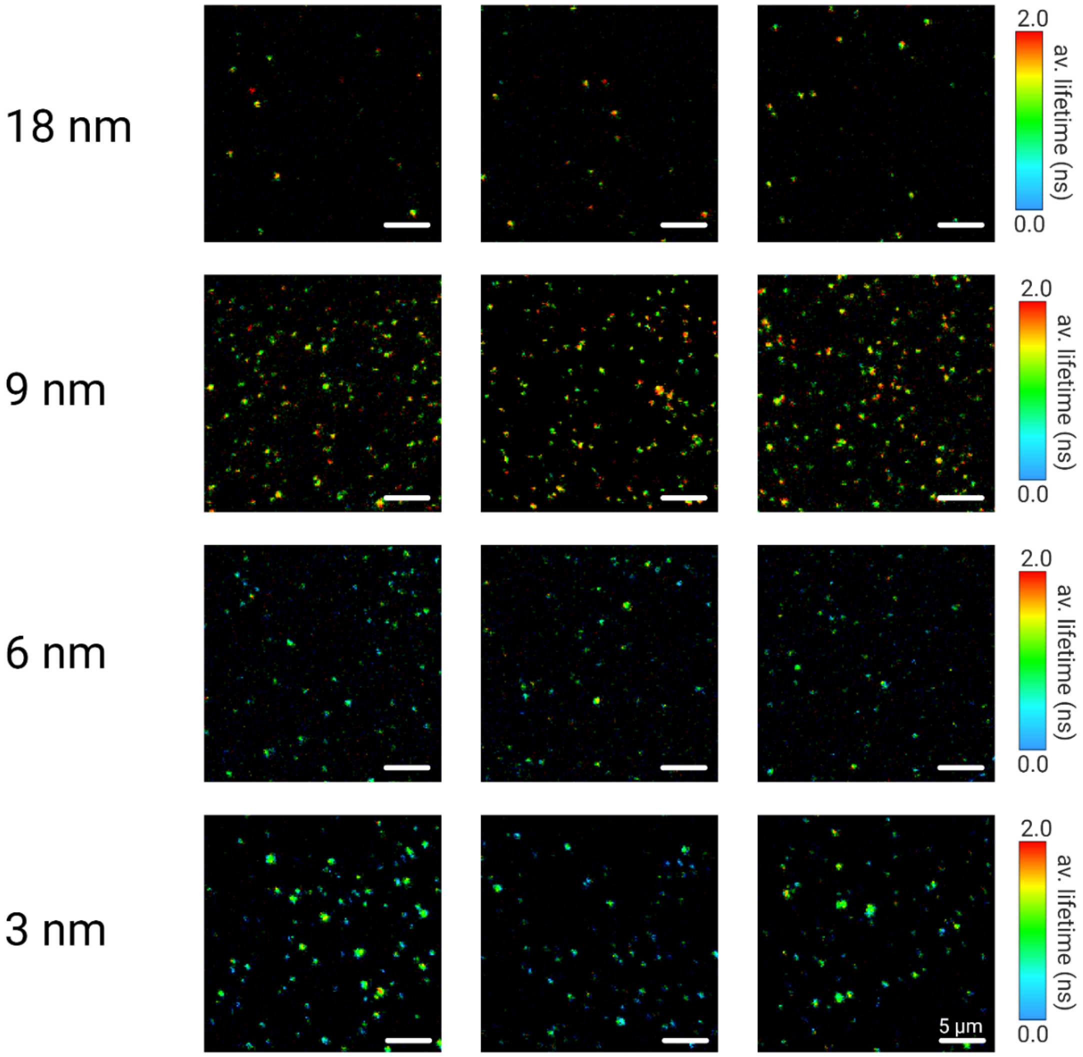


**Figure S16**. FLIM images of 18, 9, 6, 3 nm DNA origami measured in PBS, pH 7.6 containing 1 mM trolox/troloxquinone and an oxygen scavenging system measured by single-molecule sensitive confocal fluorescence microscopy excited at 640 nm with 2.5 kW $cm^{-2}$ at an integration time of 5 µs $pixel^{-1}$.

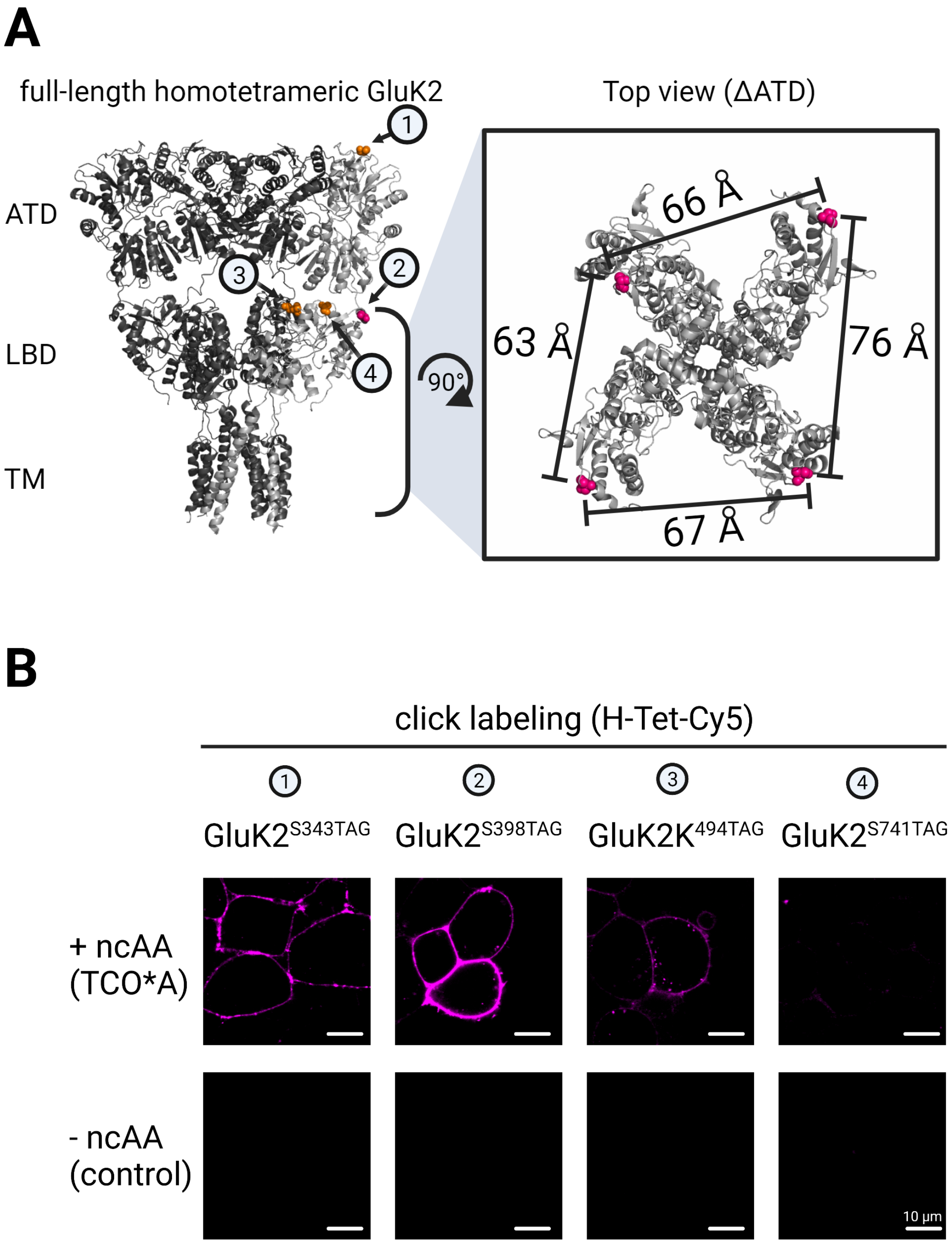


**Figure S17**. Scheme explaining click labeling and construct design of GluK2. (**A**) Different mutants 1-4 were generated within one monomeric subunit of GluK2. Calculation of the distances was performed with PyMOL (Molecular Graphics System, Version 1.2r3pre, Schrödinger, LLC) on basis of the crystal structure (PDB-ID: 5KUF). (**B**) To check the efficiency of ncAA incorporation of the different mutants, click labeling was performed with H-Tet-Cy5. Control experiments without the addition of ncAA resulted in inefficient amber suppression efficiency which leads to premature translation termination and no click labeling.

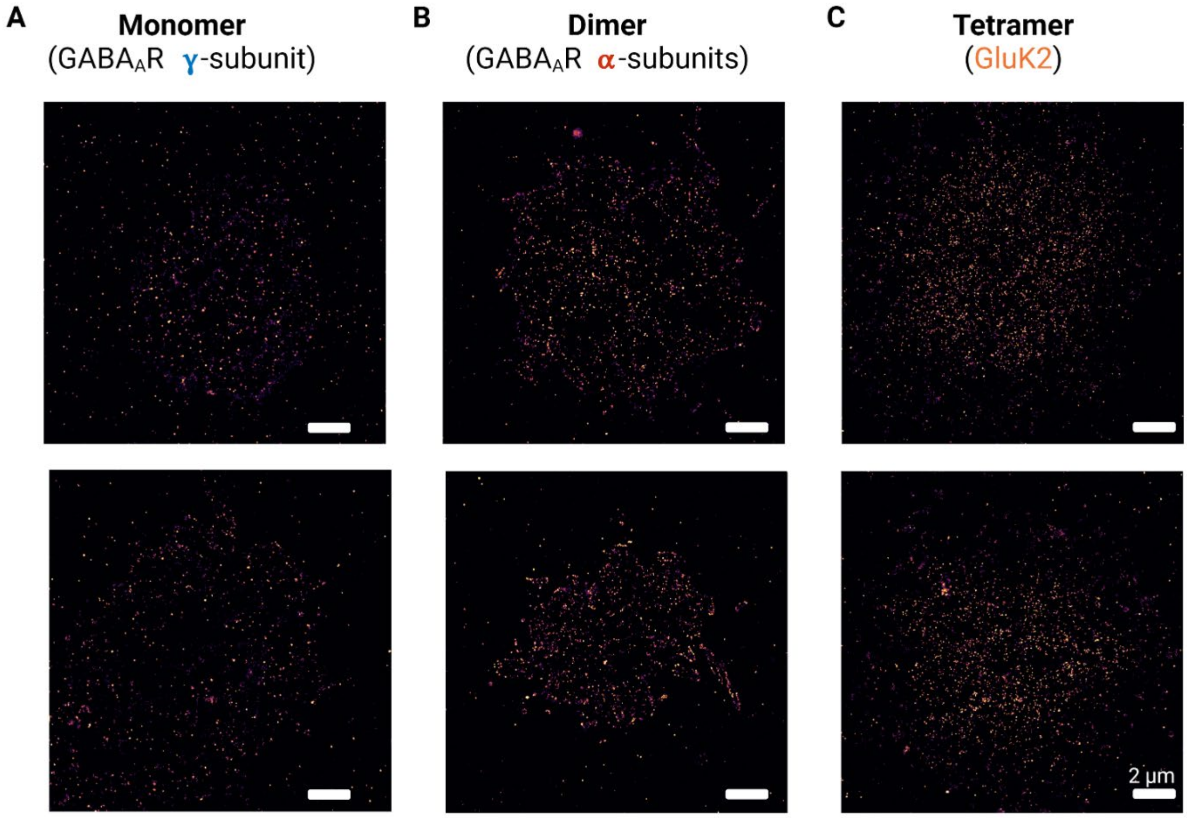


**Figure S18**. *d*STORM images of membrane receptors (20 nm pixel$^{-1}$).

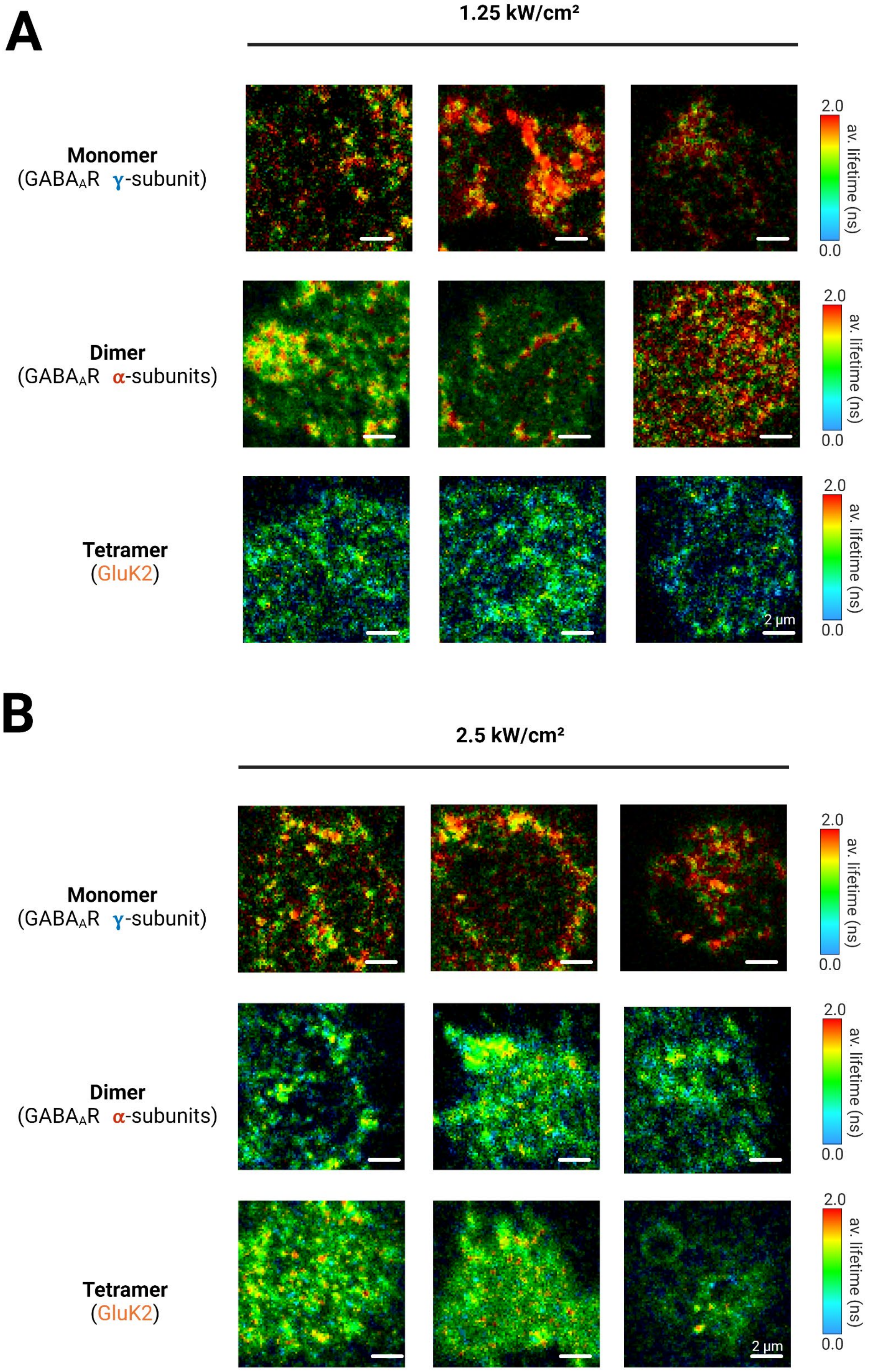


**Figure S19**. FLIM images of HEK293T cells expressing monomeric γ2-subunit of GABA-A, dimeric α2-subunit of GABA-A, and homotetrameric GluK2 receptors click-labeled with Met-Tet-Cy5 measured by confocal TCSPC imaging in photoswitching buffer at different irradiation intensity and an integration time of 5 µs $pixel^{-1}$ without applying an intensity threshold.

### SUPPLEMENTARY MOVIES

**Movies S1-S5.** DNA-PAINT movies (30 min) of DNA origami labeled with one (reference) or four docking strands separated by different distances recorded at a temporal resolution of 100 ms per frame.

**Movie S1.** DNA-PAINT of reference DNA origamis, Scale bar, 2 µm.

**Movie S2.** DNA-PAINT of 18 nm DNA origamis, Scale bar, 2 µm.

**Movie S3.** DNA-PAINT of 9 nm DNA origamis, Scale bar, 2 µm.

**Movie S4.** DNA-PAINT of 6 nm DNA origamis, Scale bar, 2 µm.

**Movie S5.** DNA-PAINT of 3 nm DNA origamis, Scale bar,

**Movies S6-S11.** *d*STORM movies (10 min) of DNA origami labeled with one (reference) or four Cy5 dyes with different interfluorophore distance recorded at a temporal resolution of 5 ms per frame.

**Movie S6.** *d*STORM of reference DNA origamis, Scale bar, 2 µm.

**Movie S7.** *d*STORM of 18 nm DNA origamis, Scale bar, 2 µm.

**Movie S8.** *d*STORM of 9 nm DNA origamis, Scale bar, 2 µm.

**Movie S9.** *d*STORM of 6 nm DNA origamis, Scale bar, 2 µm.

**Movie S10.** *d*STORM of 3 nm DNA origamis, Scale bar, 2 µm.

**Movie S11.** *d*STORM of 3 nm DNA origamis, Scale bar, 2 µm.

### REFERENCES


55. S. M. Douglas *et al.*, Self-assembly of DNA into nanoscale three-dimensional shapes. *Nature* **459**, 414-418 (2009).

56. S. M. Douglas *et al.*, Rapid prototyping of 3D DNA-origami shapes with caDNAno. *Nucleic Acids Res.* **37**, 5001-5006 (2009).

57. D. N. Kim, F. Kilchherr, H. Dietz, M. Bathe, Quantitative prediction of 3D solution shape and flexibility of nucleic acid nanostructures. *Nucleic Acids Res.* **40**, 2862-2868 (2012).

58. C. Castro, F. Kilchherr, D. N. Kim, *et al.*, A primer to scaffolded DNA origami. *Nat. Methods* **8**, 221–229 (2011).

59. V. Tretter, T. C. Jacob, J. Mukherjee, J. M. Fritschy, M. N. Pangalos, S. J. Moss, The clustering of GABA(A) receptor subtypes at inhibitory synapses is facilitated via the direct binding of receptor alpha 2 subunits to gephyrin. *J. Neurosci.* **28**, 1356-1365 (2008).

59. E.M. Petrini, T. Nieus, T. Ravasenga, F. Succol, S. Guazzi, F. Benfenati, A. Barberis, Influence of GABAAR monoliganded states on GABAergic responses. *J. Neurosci.* **31**, 1752-61 (2011).

60. A. Kuhlemann, G. Beliu, D. Janzen, E.M. Petrini,D. Taban, D.A. Helmerich, S. Doose, M. Bruno, A. Barberis, C. Villmann, M. Sauer, C.Werner, Genetic Code Expansion and Click-Chemistry Labeling to Visualize GABA-A Receptors by Super-Resolution Microscopy. *Front. Synaptic Neurosci.* **13**, 727406 (2021).

61. A. Herb, N. Burnashev, P. Werner, B. Sakmann, W. Wisden, P. H. Seeburg, The KA-2 subunit of excitatory amino acid receptors shows widespread expression in brain and forms ion channels with distantly related subunits. *Neuron* **8**, 775–785 (1992).

62. I. Nikic, et al., Debugging Eukaryotic Genetic Code Expansion for Site-Specific Click-PAINT Super-Resolution Microscopy. *Angew. Chem. Int. Ed.* **55**, 16172-16176 (2016).

63. R. Serfling, et al., Designer tRNAs for efficient incorporation of non-canonical amino acids by the pyrrolysine system in mammalian cells. *Nucleic Acids Res.* **46**, 1-10 (2018).

64. S. Wolter, *et al.*, rapidSTORM: accurate, fast open-source software for localization microscopy. *Nat. Methods* **9**, 1040–1041 (2012).

65. M. Ovesný, P. Křížek, J. Borkovec, Z. Švindrych, G. M. Hagen, ThunderSTORM: a comprehensive ImageJ plugin for PALM and STORM data analysis and super-resolution imaging. *Bioinformatics* **30**, 2389-2390 (2014).

66. K. I. Mortensen, L. S. Churchman, J. A. Spudich, H. Flyvbjerg, Optimized localization analysis for single-molecule tracking and super-resolution microscopy. *Nat. Methods* **7**, 377-381 (2010).
